# Multiscale causal network models of Alzheimer’s disease identify VGF as a key regulator of disease

**DOI:** 10.1101/458430

**Authors:** Noam D. Beckmann, Wei-Jye Lin, Minghui Wang, Ariella T. Cohain, Pei Wang, Weiping Ma, Ying-Chih Wang, Cheng Jiang, Mickael Audrain, Phillip Comella, Siddharth P. Hariharan, Gillian M. Belbin, Allan I. Levey, Nicholas T. Seyfried, Eric B. Dammer, Duc Duong, James J. Lah, Jean-Vianney Haure-Mirande, Ben Shackleton, Alexander W. Charney, Eimear Kenny, Jun Zhu, Vahram Haroutunian, Pavel Katsel, Sam Gandy, Zhidong Tu, Michelle Ehrlich, Bin Zhang, Stephen R. Salton, Eric E. Schadt

## Abstract

Though discovered over 100 years ago, the molecular foundation of sporadic Alzheimer’s disease (AD) remains elusive. To elucidate its complex nature, we constructed multiscale causal network models on a large human AD multi-omics dataset, integrating clinical features of AD, DNA variation, and gene and protein expression into probabilistic causal models that enabled detection and prioritization of high-confidence key drivers of AD, including the top predicted key driver VGF. Overexpression of neuropeptide precursor VGF in 5xFAD mice partially rescued beta-amyloid-mediated memory impairment and neuropathology. Molecular validation of network predictions downstream of VGF was achieved, with significant enrichment for homologous genes identified as differentially expressed in 5xFAD brains overexpressing VGF versus controls. Our findings support a causal and/or protective role for VGF in AD pathogenesis and progression.

**One sentence summary:** VGF protects against Alzheimer’s disease

---

Alzheimer’s disease (AD), the prevailing cause of dementia in the world, affects more than 28 million people worldwide, and its incidence is projected to double in the next 20 years (*1*). AD results in the progressive loss of cognitive function, memory, and ability to think and reason. The brains of AD patients have hallmark senile plaques in the neuropil and around brain blood vessels, composed of accumulated amyloid beta (Aβ), and neurofibrillary tangles (NFT) inside neurons, comprised of microtubule-associated hyperphosphorylated Tau protein (*2*). While therapeutic strategies to target Aβ and tau pathologies have been aggressively pursued over the past two decades, the failure to date to deliver efficacious treatments from these efforts has increased the urgency to identify and pursue different mechanisms underlying AD, such as the immune system, through microglial cells, that has more recently been shown to play a key role in AD (*3*–*9*). Furthermore, with a handful of drugs lessening some of the symptoms of AD, no effective drugs are currently available that prevent, halt or reverse the onset or progression of this disease.

One class of approaches that have delivered novel insights into the causes of AD are genome-wide association studies (GWAS), which have resulted in the identification of more than 20 risk loci falling mainly in non-coding regions of the genome (*10*, *11*), revealing a complex neurobiology with no single genetic cause. However, for most AD risk loci, the target gene or genes are difficult to identify and validate, the pathways in which these target genes operate to impact AD are largely unknown, and the broader context in which the genes and corresponding pathways relating to AD interact and the networks they form, remain largely uncharacterized. Integrative biology approaches that combine large-scale, high-dimensional data (such as DNA variation and gene and protein expression) generated in disease and control cohorts, can well complement GWAS-like approaches by employing advanced computational modeling techniques that incorporate multiple levels of data to construct probabilistic causal models of disease (or wellness) that in turn enable distinguishing between molecular traits that are simply correlated with disease, from those that are causally related (*4*, *5*, *12*–*16*). The power to infer causal relationships from large-scale data can be enhanced by systematically incorporating DNA-based variations such as expression quantitative trait loci (eQTL), as a systematic perturbation source (*4*, *5*, *13*, *15*, *17*–*34*). By integrating DNA variation data with additional types of molecular and clinical data, more complex, holistic models of disease can be constructed and then mined to elucidate regulatory and mechanistic drivers of disease, which in turn can lead to novel therapeutic points of intervention.

Here, we employed probabilistic causal reasoning to organize DNA, RNA, protein, and clinical data we and others have generated as part of the Accelerating Medicines Partnership-Alzheimer’s Disease (AMP-AD; https://www.synapse.org/#!Synapse:syn2580853/wiki/409840) on a population of late-onset AD individuals and controls, to construct a predictive network model of AD, providing a comprehensive characterization of the complex architecture of AD in the human brain. Because the networks that result from this process represent different scales of data (DNA, RNA, and protein expression), we refer to them as multiscale network models of disease. Given these multiscale networks, the causal links among the nodes comprising them can be mined to identify gene or protein expression traits that are predicted to modulate network states that in turn drive AD. The identification of these master drivers of the disease networks provides an objective, data driven way to uncover novel causal regulators of disease. Strikingly, among the key driver (KD) genes we identified was VGF, a nerve growth factor (NGF) and brain-derived neurotrophic factor (BDNF) inducible gene. VGF encodes a protein and neuropeptide precursor the actions of which are in part BDNF/TrkB-dependent (*35*, *36*).

Although VGF has been reported to regulate fear and spatial memories in mouse models ((*35*, *37*, *38*), and to be an AD biomarker, with VGF-derived peptides found to be reduced in the CSF of AD patients compared to healthy controls (*39*–*46*), VGF has not previously been causally associated with AD. We determined through our network models that VGF was the only downregulated KD for AD that was conserved across the RNA, protein, and combined RNA and protein networks we constructed. We replicated these findings in other brain regions (*47*) and in an independent dataset (*48*, *49*), and observed evidence of genetic association in the largest AD GWAS to date (*10*). Given VGF’s status as the top KD we identified in our networks, we overexpressed VGF in the 5xFAD mouse model of familial AD and found that it not only lowered overall amyloid plaque and Tau-associated dystrophic neurite levels, but it significantly perturbed gene expression traits that were enriched for genes predicted by our networks to change in response to VGF modulation. Taken together, these results provide molecular and functional validation of our multiscale causal network analysis finding of *VGF* as a driver of AD pathophysiology. We conclude that the genes and clinical features linked to *VGF* provide novel insights into the mechanisms underlying AD risk and pathogenesis.

## Results

Our overall strategy for elucidating the complexity of AD is depicted in Fig. 1 (Fig. S1) and is centered on the objective, data-driven construction of predictive network models of AD that can then be queried to identify network components causally associated with AD. The master regulators that modulate the state of these AD-associated network components can then be readily identified from the network model. We have previously developed and applied the network reconstruction algorithm, RIMBANET, which statistically infers causal relationships between DNA variation, gene expression, protein expression and clinical features that are scored in hundreds of individuals or more (*5*, *13*–*15*, *18*, *19*, *27*, *29*, *32*, *34*, *50*, *51*).The inputs required for this type of analysis are the molecular and clinical data generated in populations of individuals, as well as first order relationships between these data, such as QTL mapped for the molecular traits and causal relationships among traits inferred by causal mediation analysis that uses the mapped QTL as a source of perturbation. These relationships are input as structure priors to the network construction algorithm, boosting the power to infer causal relationships at the network level, as we and others have previously shown (*5*, *13*, *15*, *25*, *27*, *32*–*34*, *51*).

**Fig. 1.**
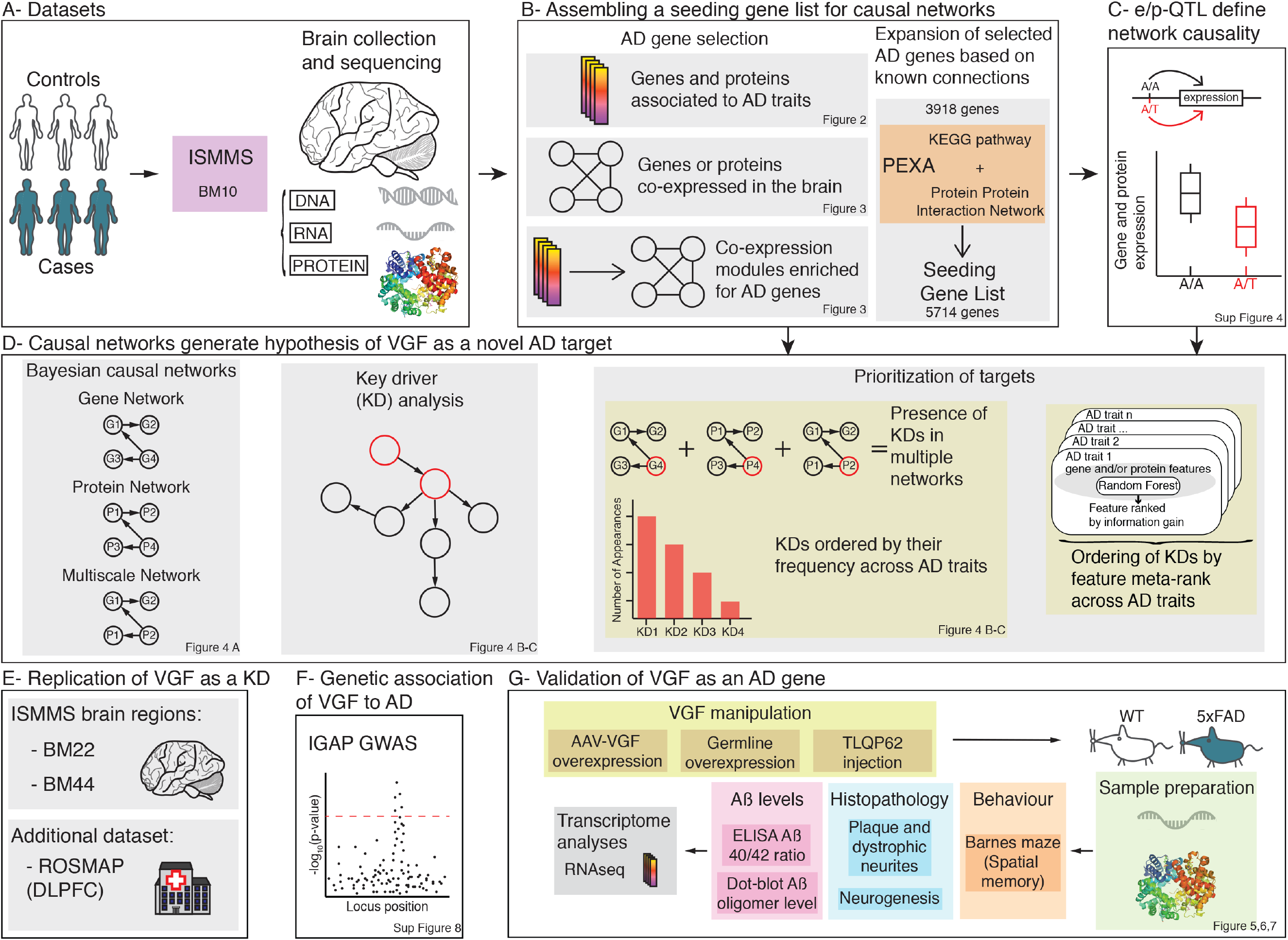
Pipeline Overview. Description of data followed by schematics highlighting analyses and validation workflows performed to identify and validate targets of AD, as described in the results.

In this study, the input data to construct predictive network models of AD were generated as part of the AMP-AD consortium, and included whole exome sequencing (WES), RNA sequencing (RNA-seq), and protein expression data from the anterior prefrontal cortex (Brodmann area 10, BM10) in a large cohort of post-mortem samples from the Mount Sinai Brain Bank (MSBB, n=315), across the complete spectrum of AD clinical and neuropathological traits (from controls to neuropathologically-proven AD, Fig. 1A) (*47*). To focus the input of molecular traits into the network reconstruction algorithm on those traits associated with AD, we examined the association between the molecular data and AD clinical and neuropathological features, and identified AD gene and protein expression signatures comprised of hundreds of gene and protein expression traits. These signatures were enriched for mitochondrial and immune processes. To identify gene and protein expression traits co-regulated with the AD signature genes, we constructed gene and protein co-expression networks, and from these networks identified highly interconnected sets of coregulated genes (modules) that were significantly enriched for the AD expression signatures as well as for pathways previously implicated in AD (Fig. 1B). To obtain a final set of genes for input into the causal network construction process, we combined genes in the AD expression signatures and genes in the co-expression network modules enriched for these signatures (the seed set). We then expanded this seed set by incorporating prior pathway knowledge from the literature to ensure we did not miss important AD genes due to a lack of power in the differential expression analysis to identify them (Fig. 1B).

With our AD-centered input set of genes for the network constructions defined, we mapped gene and protein quantitative trait loci (eQTLs and pQTLs, respectively) for expression traits in this set to incorporate the QTL as structure priors in the network reconstructions, given they provide a systematic perturbation source that can boost the power to infer causal relationships (Fig. 1C)(*5*, *13*, *15*, *25*, *27*, *32*–*34*, *51*). The input gene set, and eQTL/pQTL data from MSBB were then processed by RIMBANET to construct probabilistic causal networks of AD (Fig. 1D). An artificial intelligence algorithm to detect key driver genes from these network structures was then applied to identify and prioritize master regulators of the AD networks (Fig. 1D). Our findings were then replicated in other datasets (Fig. 1E), and were supported by genetic associations to AD in the largest GWAS to date (Fig. 1F). For the top regulator we identified, VGF, we performed functional and molecular validation in the 5xFAD mouse model (Fig. 1G).

### The Mount Sinai Brain Bank Study Population and Data Quality Control

The AD and control populations profiled in this study are a component of the MSBB (*47*). From the > 1,900 participants making up this brain bank, 117 definitive AD cases were selected for this study, along with 123 possible and probable AD cases and 75 non-demented controls. The selection criteria were neuropathological evidence of AD by CERAD (*52*) classification or no neuropathological evidence of AD. In addition, donors with neuropsychiatric disease and/or comorbid neurodegenerative diseases, and/or neuropathologically significant cerebrovascular disease, were excluded. A summary of the MSBB population demographics is provided in Data S1. DNA, RNA, and protein were isolated from the BM10 region of study participants for molecular profiling (Fig. 1A).

The DNA and RNA sequencing data were processed using standard pipelines, including quantification of gene expression, variant detection and QC for the RNA-seq data (*53*) (see methods and materials for details). We identified 18 samples for which variants identified from the RNA-seq and DNA WES data did not achieve the level of concordance expected for samples derived from the same donor. In addition, for 6 samples the sex inferred by the DNA and RNA data did not match the sex reported for the corresponding participant in the clinical report. Finally, 13 of the RNA-seq samples mapped to more than one WES sample (the discordance rate with the best matching sample was > 10%). We removed from all further analyses 16 of these samples that could not be unambiguously corrected (Fig. S2A, S2B, Data S1), leaving 279 for detailed analyses.

To assess the integrity of these data and identify covariates that could impact our analyses, we carried out variance partition (*54*) and principal component (PC) analyses, and identified exonic mapping rate (fraction of reads mapping to exonic regions), RNA integrity number (RIN), and sequencing batch as covariates explaining the greatest variation in gene expression across samples. In order to minimize the impact of these covariates on detecting our primary signal of interest (association of molecular traits to AD), we adjusted the normalized RNA-seq count data by correcting for race, sex and the main drivers of technical variation using PC analysis (Fig. S2C, S2D, S2E, S2F) and linear mixed models, which included post-mortem interval (PMI), RIN and exonic mapping rate. Protein expression data were processed in a similar fashion and corrected for batch, PMI, race and sex to minimize unwanted variation.

### Identifying an AD-centered gene set to construct a predictive model of AD

To construct the AD-centered predictive network models, we constrained the number of inputs into the reconstruction process to those supported by the MSBB data as associating with AD, with this reduction in dimensionality also providing a more computational tractable path for the network constructions. Our first step in this process was to identify gene and protein expression traits associated with AD (Fig. 1B). To cast the most comprehensive net for AD-associated features, we first examined the association between the molecular expression traits and clinical/neuropathological features used to characterize AD. Given the complexity of AD, 6 clinical and neuropathologic characteristics were used to define the severity of disease in patients, including clinical staging with the clinical dementia rating (CDR), pathological staging of neurofibrillary tangles or Braak score (bbscore), clinical neuropathology diagnosis (PATH.Dx), CERAD neuropath criteria (CERJ), neuropathology category (NP-1) and mean cortical neuritic plaque density (PlaqueMean). We characterized the differences and similarities specific to each of these disease traits by examining their canonical correlation structure with one another in the MSBB population (Fig. 2A). While these cognitive and neuropathological measures of AD were highly correlated, visible variation among them highlights their complementary nature, with non-overlapping signals that may represent different aspects or subtypes of AD. Thus, we constructed DE signatures for each of these clinical AD features.

**Fig. 2.**
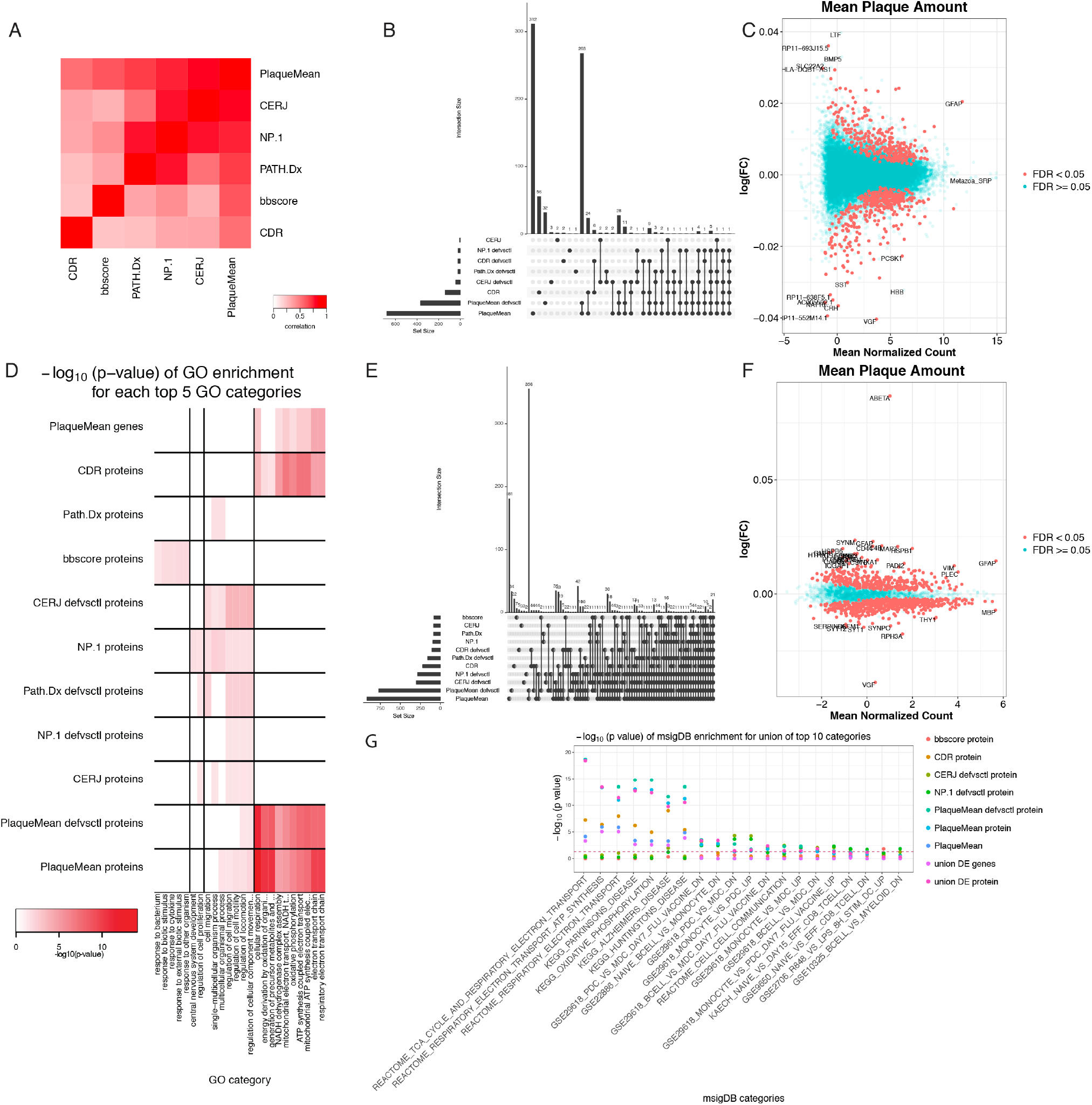
Characterization of AD traits and brain gene and protein expression. **A)** Canonical correlation heatmap of disease traits. The intensity of the red color indicates the strength of the correlation between traits. The x and y axes represent the traits: clinical dementia rating (CDR), Braak score (bbscore), clinical neuropathology (PATH.Dx), neuropathology category (NP.1), CERAD neuropath Criteria (CERJ), mean neocortical plaque density (number of plaques/mm^2^, PlaqueMean). **B and E)** Breakdown of DE genes **(B)** and proteins **(E)**. Shown is the UpsetR plot (methods) of the DE genes or proteins overlapping across tests. The bars represent the set sizes and the points which category the set size represents. **C and F)** PlaqueMean DE genes **(C)** and proteins **(F)**. The x and y axes are the mean normalized count for each gene or protein and their log fold change. Blue and red genes and proteins are representative of FDR larger and smaller than 0.05, respectively. The strongest DE genes are highlighted on the plot. **D)** GO term enrichment across all signatures. The heatmap depicted represents the −log10(FDR) of the top 5 significant GO terms associated to signatures across all traits. Rows are GO terms and columns signatures. **G)** MsigDB pathway enrichment for all signatures. The barplot represents the union of the top 10 significant MsigDB categories associated to signatures for all traits; x and y axes are MsigDB terms and the −log10(FDR); colors represent the traits.

As defined in Table S1, we computed differential expression (DE) signatures for AD by comparing controls against individuals with any level of dementia or pathology, and then controls against individuals with neuropathologically-proven AD (definite AD). In this way, we generate signatures across the range of disease. We detected significant DE signatures at a false discovery rate (FDR) < 0.05 for most traits across the disease spectrum (Fig. 2B, Fig. 1B). The PlaqueMean feature generated the largest DE signature (Fig. 2C, 2D), with the Gene Ontology (GO) term ‘respiratory electron transport chain’ (fold enrichment, FE, = 4.9, FDR = 4.42e-5) identified as the most enriched pathway. From the log fold-change, log(FC), distribution (Fig. 2C) of the PlaqueMean signature, the gene VGF, nerve growth factor inducible, is identified as the gene with the largest negative log(FC) (more highly expressed in controls than cases). VGF was previously shown to be downregulated in patients with familial AD (*55*), which is consistent with our findings here. The DE signatures for the other disease traits (Data S2) are depicted in Fig. S3A.

We ran a similar DE analysis to identify AD signatures from the protein expression data (Fig. 2E, Fig. 1B, Fig. S3B), and found that significant DE protein signatures were identified for all AD clinical features, with PlaqueMean again giving rise to the most significant signature (Fig. 2E). For each clinical or neuropathological trait, the protein with the highest log(FC) was Aβ, followed by other known AD proteins such as MAPT, GFAP, HSPB1, RPH3A, SYT1 and PADI2 (*56*–*60*). Strikingly, as with the gene DE signature, the protein with the lowest log(FC) was VGF, highlighting the strong dysregulation of the gene/protein product in AD brains (Fig. 2F). Several protein DE signatures were enriched for GO terms (Fig. 2D), with the GO term ‘cellular respiration’ as the most significant in the PlaqueMean protein DE signature (FDR = 8.3e-15, FE = 2.4). The electron transport chain pathway and AD KEGG pathway from the MsigDB and KEGG databases, respectively, were also significantly enriched in the PlaqueMean protein DE signature (Fig. 2G, Data S3).

From the DE analysis we took the union of all genes across all signatures to form a preliminary set of AD-associated input features for the network reconstructions. This set was comprised of 788 genes identified from the gene expression signatures and 1016 genes from the protein expression signatures at an FDR < 0.05, with 55 genes overlapping, demonstrating the highly complementary nature of the gene and protein expression data. These resulting sets of 788 genes and 1018 proteins are referred to as the AD DE signature sets.

DE analysis provides the most straightforward way to uncover patterns of expression that associate with AD. However, the power in such an analysis is limited with respect to expression differences that are moderate to small. To complement DE analysis to identify AD-associated genes, we clustered the genes and proteins into data-driven, meaningful functional biological groups by constructing co-expression networks, which have enhanced power to identify coregulated sets of genes (modules) that are likely to be involved in common biological processes. Co-expression modules that are enriched for genes associated with AD implicate all genes in the module as potentially AD-associated, even if they were not identified as DE.

The gene co-expression network was comprised of 24,865 genes and 29 modules (Fig. 1B, Data S5), while the protein co-expression network consisted of 2,692 proteins organized into 9 modules (Fig. 1B, Data S4), with most modules (26 and 8 gene and protein modules respectively) having significant GO term associations at an FDR < 0.05 (Fig. 3A, 3C). To assess which sets of modules were associated with AD, we projected the DE signature set onto the coexpression network modules (Fig. 1B, 3B, 3D). We identified 4 modules from the gene coexpression network that were significantly enriched for the gene AD DE signature set (Fig. 3B). These modules were enriched for the GO terms ‘induction of positive chemotaxis’ (greenyellow, FDR = 3.0e-2), ‘histone modification’ (peru, FDR = 1.7e-3), ‘mitochondrion organization’ (pink, FDR = 1.9e-5) and ‘synaptic transmission’ (yellow, FDR = 1.6e-5) (Data S4). For the protein coexpression network, we identified 3 protein modules as enriched for the protein AD DE signature set (Fig. 3D). These modules were enriched for ‘synaptic transmission’ (blue, FDR = 4.6e-15), ‘response to molecule of bacterial origin’ (green, FDR = 5.9e-3) and ‘energy derivation by oxidation of organic compounds’ (yellow, FDR = 2.8e-14) (Data S4).

**Fig. 3.**
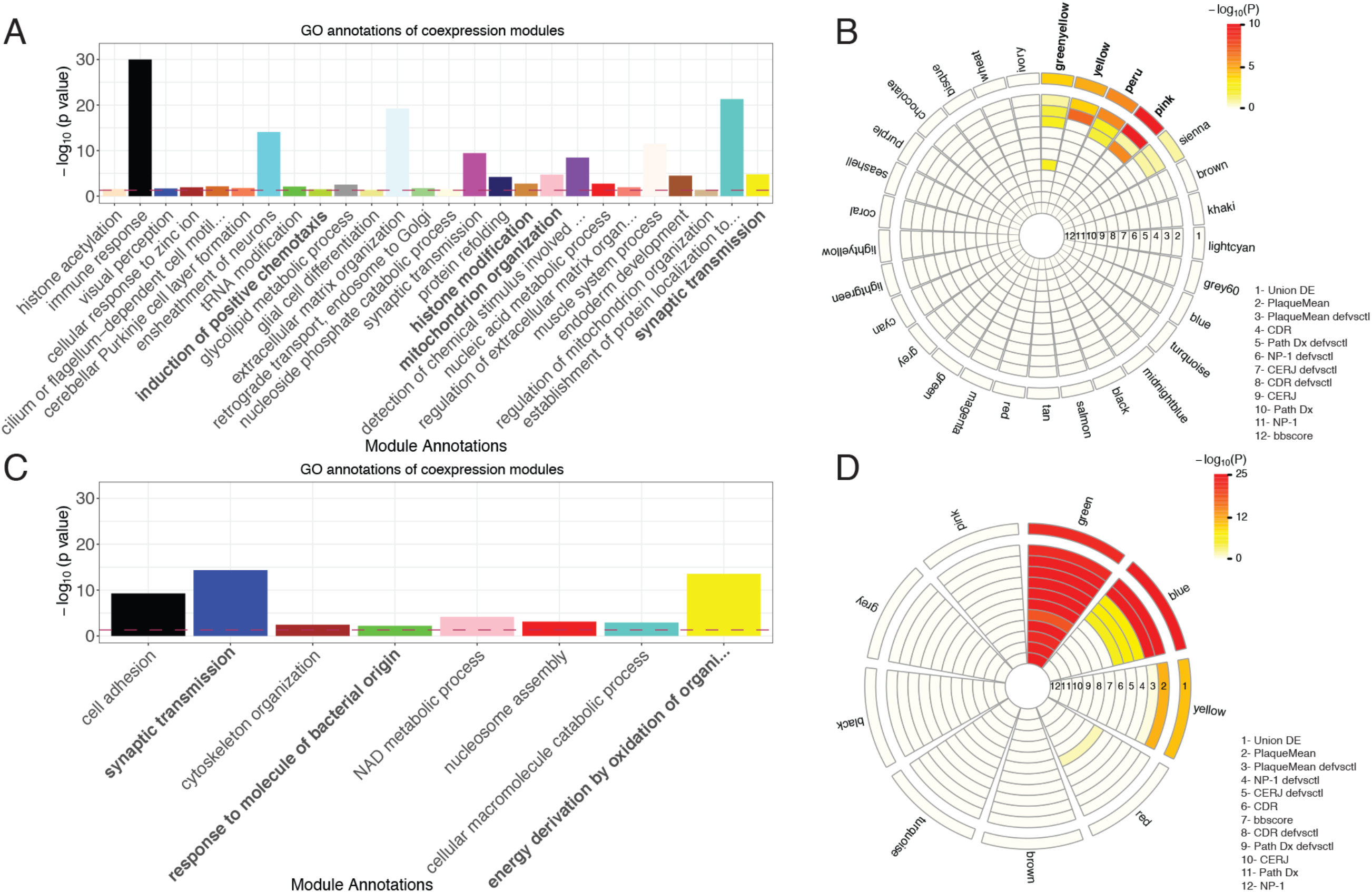
Co-expression analyses. **A and C)** Top GO annotations for gene **(A)** and protein **(C)** coexpression modules. The x and y axis represent the best GO term associated with each module and the −log10(p value) of the enrichment, respectively. The color of the bars represents the module names. In bold are the 4 modules enriched for genes in the union of the DE signatures. **B and D)** Module enrichments for DE genes **(B)** and proteins **(D)**. The circos plot depicts the enrichment of each module for DE genes or proteins; the “hotness” of the color represents the magnitude of the −log10(p value) of the enrichment for the corresponding signature list. The traits are defined as 1 through 12; in bold are the modules enriched for the union of the DE signatures.

The construction of co-expression networks from the combined gene and protein expression traits resulted in modules comprised nearly exclusively of one type of data (either gene or protein expression) (Data S4). While technical components of variation specific to technologies used to score the gene and protein expression will partly explain this pattern of coexpression, given traits of a particular type are more correlated to traits of that same type than traits of other types, the complementarity of the gene and protein expression plays a role as well. For example, while RNA measures generally reflect expression levels in cells local to the brain region assayed, select RNAs or RNA isoforms that are known to be transported into dendrites (e.g. BDNF long 3’ UTR mRNA) could potentially contribute to this signal as well (*61*–*63*). Similarly, protein measures may reflect proteins synthesized in the local brain region that was profiled, proteins that are transported in secretory vesicles via neural pathways from cell bodies in distal regions, and proteins that are locally translated from mRNAs transported from distal regions. Thus, simultaneous sampling of RNA and protein expression in a specific brain region provides complementary data sets that not only reflect linear DNA to RNA to protein synthesis, but that also capture dynamic changes in the flux of transported proteins and RNAs into the local region. Co-expression networks primarily capture linear relationships among traits, but largely ignore the nonlinear relationships that exist between gene and protein expression levels. As seen below, Bayesian networks address this issue given their ability to capture nonlinear interactions.

We therefore expanded the DE signature set of input genes for the predictive network constructions by taking the union of the DE signature set and all genes across all gene coexpression modules enriched for genes in the DE signature set, resulting in a set of 3,918 genes, referred to here as the expanded DE signature set (Fig. 1 B).

### Genetic modulation of gene and protein expression in the prefrontal cortex

Central to our approach to construct predictive network models is the integration of QTLs as a systematic source of perturbation to enhance causal inference among molecular traits, an approach we and others have demonstrated across a broad range of diseases and data types (*5*, *13*–*15*, *17*, *19*, *21*, *24*, *25*, *27*, *28*, *30*, *31*, *33*, *34*, *51*, *64*–*71*). QTL mapping identifies DNA loci that associate with quantitative traits such as gene and protein expression, aiding in the identification of regulatory and mechanistic relationships among genes and proteins, which in turn can provide critical insights into biological processes related to the functioning of cells and their association to disease. Given we scored gene and protein expression traits in this study, we mapped eQTL and pQTL for all molecular traits to identify significant QTL as inputs along with the AD-associated expression traits into the network reconstruction process.

We found 4,224 genes with at least one eQTL (eGenes) and 158 proteins with at least one pQTL (eProteins), at a 0.05 FDR (Data S5). To assess the degree of conservation across the RNA and protein domains and to help illuminate AD genetics, we characterized the number of QTLs overlapping the expanded AD DE signatures (Fig. 1C, Fig. S4A). Of these, we identified 83 proteins with pQTLs, 683 genes with eQTLs, 7 genes and proteins with both e-and pQTLs. Fig. S4B shows an example of a gene, *GSTM3*, whose cortical gene and protein expression levels are associated with a shared SNP, rs1332018 (*72*). Given the relationship between transcripts and the proteins they encode, we applied a causal mediation test (*25*, *73*) to assess whether changes in gene expression induced by the eQTL were causal for the corresponding changes in protein expression for the 33 product pairs under control of the same SNP. Interestingly, the causal mediation test supported 26 products of gene and protein expression as being independently regulated by the *cis* variation (Data S5), suggesting post-translational events may be impacted by the same *cis* variation that impacts transcription, albeit in an independent fashion, perhaps partially explaining the low correlation we and others have observed between gene and protein expression (as highlighted below).

Of the eQTLs and pQTLs identified, 766 corresponded to genes and proteins overlapping the expanded AD DE signature and so were included as inputs into the network constructions.

### Identification and prioritization of key driver genes identified from predictive network models of AD

To elucidate the structure of the complex interactions represented in the expanded AD DE signature set and associated QTL, we employed a Bayesian network (BN) modeling approach (Fig. 1D). BNs are graphical models that capture relationships (depicted as edges) among nodes (gene or protein expression traits) systematically across high-dimensional datasets. BNs not only capture linear correlations and higher order correlations among nodes (like co-expression networks), but can also capture nonlinear relationships and infer causal links that define information flow, thereby providing a richer, more informative context for discovery (*5*, *15*, *18*, *25*, *32*, *34*, *51*). Because the number of possible networks to search to identify the network that best fits the data grows exponentially with the number of nodes in the network, a brute force search of all networks to find the best one is not feasible (*51*). Heuristics are used to constrain the size of the search space to efficiently search it (*34*). Towards that end, we constructed the AD-focused seeding gene set to reduce the size of the search space, with the core of this set comprised of the AD DE signatures (Data S2) that was then expanded to include all genes in the co-expression network modules significantly enriched for these core AD signature genes (Data S4).

However, a limitation of this empirically determined gene set is that it may miss important genes due to nonlinear interactions not captured by co-expression networks, a lack of power to detect all relevant genes in the gene expression data, or genes active in tissues or stages of disease that were not as well captured in the MSBB population. To account for this, we further expanded the seeding gene set (3,918 genes) using the PEXA algorithm (*74*), which enables inclusion of genes from literature-derived pathways that are linked to the core genes or genes that interacted with coding products of the core gene set in protein-protein interaction (PPI) networks. The application of PEXA resulted in the identification of an additional 1,796 genes, bringing our final list of genes to use in the construction of the BNs to 5,714 genes (Fig. 1B, Data S5), compared to the 24,865 transcripts detected as expressed in the MSBB dataset.

From the seeding gene set we constructed 3 BNs, one for each data type and one multiscale gene and protein expression BN (Fig. 1D, Fig. 4A, Data S4). The BN construction process integrated QTLs as structure priors to both reduce the size of the search space and enhance causal inference among nodes (*25*, *34*). For the gene-only and the multiscale networks, we included all genes in the seeding gene list described above. However, given that the number of protein expression traits was smaller than the reduced gene expression-based set, we included all 2,692 proteins detected in the protein expression set in both the protein-only and multiscale networks. Finally, to further constrain the search space for the multiscale network construction, we added a second weaker structure prior that increased the likelihood for edges corresponding to genes and the proteins for which they code (motivated by the central dogma of biology).

**Fig. 4.**
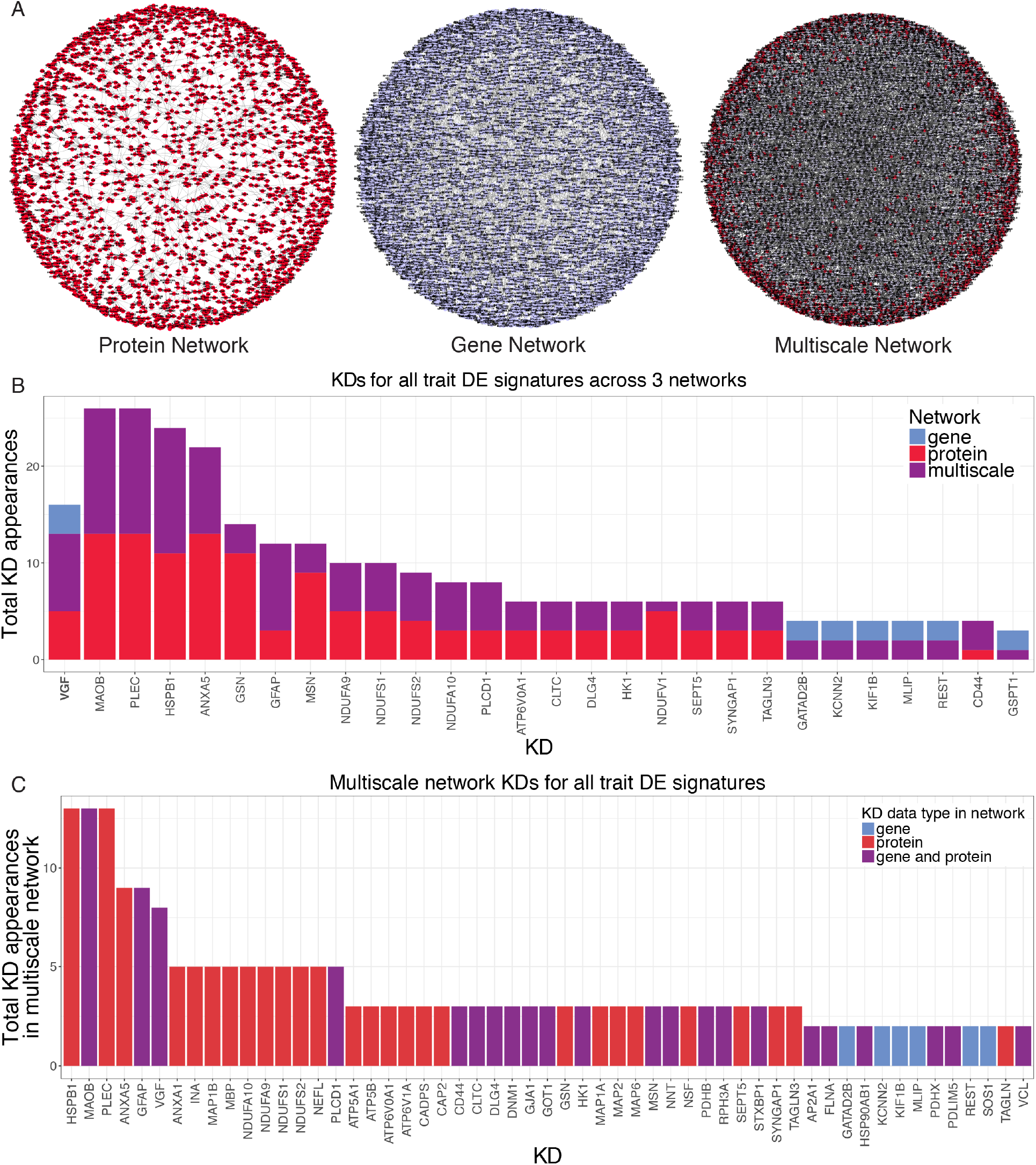
Bayesian Causal Networks and Key Drivers. **A)** Bayesian networks. Visualization of the 3 AD networks described in the main text using an edge-weighted spring embedded layout. The red nodes are proteins and the blue nodes genes. **B)** Key driver (KD) genes and proteins of the DE signatures across all 3 networks. The x and y axes depict the different KDs appearing in at least 2 networks and the number of times they are identified as KDs for DE signatures across all 3 networks. The colors of the bars are indicative of the network of origin of the KDs. **C)** KD of DE signatures in the multiscale network, as described for panel B. The color of the bars is indicative of KDs presence only in the gene expression, protein expression or in both.

While including such structure priors as a guide can result in more accurate networks, such priors are not absolute for any prior edge, but rather, any such edge must ultimately be supported by the data, an important feature given our finding from the causal mediation analysis described earlier that did not always support causal relationships between gene and protein expression traits.

Given the resulting BNs infer the causal flow of information, they can be queried to identify major points of regulatory control. Thus, we analyzed each of the BNs to identify master causal regulators (referred to as key driver genes/proteins, or KD) that are predicted by the network to modulate its state. Key Driver Analysis (KDA) (*75*) was developed for this purpose to identify nodes in the network that are predicted to modulate a significantly enriched proportion of nodes comprising a subnetwork of interest. In the context of AD, KDs of interest are those predicted to modulate components of the network that are enriched for gene and protein AD DE signatures. Thus, to predict AD KDs, we projected the DE signatures for each AD clinical and neuropathological trait onto each of the 3 BNs. Each network projection consisted of overlapping gene and protein nodes from the DE signatures with all nodes in their respective BNs. We further extracted all nodes in each network within a path length of 6 (layers) in this overlap, and identified the largest connected graph from this set of nodes and all associated edges. KDA was then carried out on each of the subnetworks resulting from these projections, resulting in a list of 499 unique KDs at FDR < 0.05 across the 3 networks (Fig. 1D, Data S6).

We next characterized the distribution of counts regarding the number of times a gene or protein was identified as a KD across all projections (Fig. 4B; Fig. 1D). The multiscale network structure was of particular interest, given it resulted from the integration of gene and protein expression data in the MSBB population (Fig. 4A). The KDs identified from this single coherent network structure are depicted in Fig. 4C. Only one KD, *VGF,* was found to be conserved across all 3 networks, supporting its potential importance in AD. Other KDs that appeared in multiple networks included genes already known to be important for AD, including GFAP, MAOB and GSN (*58*, *76*, *77*).

To complement prioritizing the importance of a KD by its replication across multiple networks as a key driver gene, we rank-ordered all KDs with respect to their importance in classifying AD cases and controls. We constructed 500 random forest classifiers for each of the 11 AD clinical and neuropathological features to distinguish AD cases from controls, as defined by these clinical features, for each scale of data separately as well as combined. All protein and gene expression traits were used to construct the classifiers, and all traits were rank-ordered based on a weighted z-score measure that combined trait-specific information scores across all classifiers, which indicates the explanatory power a given trait has to distinguish AD cases from controls, weighted by the prediction power of that classifier (area under the curve) (Data S7, see methods). VGF was again consistently identified in the top ranked KDs that had not previously been causally implicated in AD (gene network rank: 10, protein network rank: 2, multiscale network rank: 2) (Data S6).

Given *VGF* was the only KD gene identified across all AD networks, the top ranked KD gene in the AD classifiers that had not previously been causally associated with AD (VGF was ranked second overall), and was the top upregulated KD gene in controls (predicted to be the most protective against AD) in the multiscale network, we pursued VGF for extensive experimental validation (Fig. 1G). The causal networks identifying VGF also provide a context that can aid in understanding the mechanisms of action for genes such as *VGF*. If we identify the subnetwork across all three MSBB AD BNs comprised of nodes within a path length of 2 of *VGF* in each network (Fig. 7A), we see that Aβ and other known AD genes, such as HSPB1, CLU, MAOB, RPH3A, *FOSB*, and *BDNF* (*56*, *60*, *77*–*80*) are either directly connected to *VGF* or only one path length away.

### Validation of VGF as a Key Driver gene in AD

To validate VGFs role as a key driver gene of AD, we attempted to replicate our finding across different brain regions and in independent datasets, we examined VGF for association to AD in genetic studies, and we directly tested VGF in an experimental model of AD to test prospectively our prediction of VGF as playing a causal and/or protective role in AD pathogenesis and progression.

#### VGF replicates across different brain regions and in independent datasets

To further support VGF as a KD for AD and assess its regulatory role in different brain regions, we applied the same analysis pipeline (defined in Fig. 1A–1D), allowing for slight variations required to adapt our process to these data, to multiple brain regions in the AMP-AD MSBB dataset. We identified VGF as a KD in two of the three additional brain regions in the MSBB dataset, the superior temporal gyrus (BM22, Data S8) and the pars opercularis (BM44, Data S9). VGF did not reproduce as a KD in the brain region that seemed to be most affected by the disease (highest number of DE genes), the ectorhinal area (BM36, Data S10), potentially reflecting a complete disruption of the regulatory network in regions of the brain already badly damaged by the disease. We further applied the analysis pipeline defined in fig. 1A–1D on a completely independent dataset, the ROSMAP dataset (*48*, *49*), using DNA and RNA data generated in the same brain region as our original result, the dorsolateral prefrontal cortex (PFC). In the ROSMAP PFC network that resulted, VGF was again identified as a key driver gene (ROSMAP, Data S11).

#### Genetic support for VGF association to AD

DNA variations in and around VGF have not previously been identified as associating with AD. In addition, we did not identify any e‐ or pQTL for VGF, although one of the strengths of a more integrative, causal network-based approach is the ability to infer causality in a complementary way to more direct methods such as genome-wide association studies. However, to assess whether there is genetic support for VGF associating with AD given larger genomic datasets, we tested DNA variations in the VGF gene region for association to AD in the largest AD GWAS to date (*10*). We defined a 125KB region around VGF and identified a peak characteristic of an AD-associated region, with the most significantly associated SNP in the region having a p = 0.00004, which was significant even after Bonferroni correction for the 727 SNPs tested in the region (corrected p = 6.8e-5) (Fig. S5). While such results are not genome-wide significant, given we targeted a single gene region with a specific hypothesis, the results do provide support for an association.

#### In vivo molecular and physiologic validation of VGF as an AD key driver gene

To directly validate the role of VGF as a KD gene in AD pathogenesis and progression (Fig. 1F), we modulated VGF levels in the transgenic 5xFAD amyloidopathy mouse model that express human PS1 and APP containing five familial AD mutations (*81*). 5xFAD mice were crossbred to a VGF germline knock-in mouse model (VGF^Δ/Δ^ (*82*)), in which insertion of a pgk-neo selection cassette into the *Vgf* 3’UTR leads to a VGF mRNA truncation in the 3’UTR region (Δ3’UTR), resulting in increased protein translation and elevated VGF protein levels in mouse brain (Fig. S6A).

Levels of VGF protein in VGF^Δ/Δ^ hippocampus are modulated in a similar ‘physiological range’ (increased to ~150-200% control) as VGF protein levels are altered in male mice following chronic social defeat stress (decreased to ~50% control in hippocampus and increased to ~140% control in nucleus accumbens). VGF mRNA levels are similarly regulated in human control subjects and patients with major depressive disorder (MDD), being reduced in hippocampus to ~50% control in male and female MDD patients and increased in male MDD nucleus accumbens to ~150% control (*83*).

Using VGF^Δ/Δ^ mice to increase germline VGF expression in this physiological range, we assessed whether elevated levels of VGF modulate Aβ deposition in the brain of 10-month-old 5xFAD mouse, quantifying Aβ deposition by immunohistochemistry using 6E10 antibody. We found a dramatic decrease in 6E10 immunoreactive plaques in both cortical and hippocampal regions of 5xFAD,VGF^Δ/Δ^, compared to 5xFAD brains, while total brain transgenic APP protein levels remained unchanged (Fig. 5A, Fig. S6B). Microglial activation in AD patients (*84*–*87*) and increased microglial number and in some cases activation in 5xFAD (*81*, *86*, *88*) have been reported, suggesting a pathological connection between amyloid deposition and neuroinflammation. The number of Iba-1-positive cells, a microglial marker, was also significantly reduced in the cortex of 5xFAD,VGF^Δ/Δ^ compared to 5xFAD with normal levels of VGF (Fig. 5A). In addition, rapid and aggressive amyloid pathology in 5xFAD was associated with reduced neuron numbers and neurogenesis in the subgranular zone of hippocampus (*89*), which was fully rescued by VGF germline overexpression (Fig. 5B). Increased levels of Tau phosphorylation (p-Tau) are observed in the clusters formed by dystrophic neurites around amyloid plaques in the brains of human patients and mouse AD models (*90*, *91*), including 5xFAD (Fig. S6C). These were reduced by germline VGF overexpression in 5xFAD,VGF^Δ/Δ^ mice (Fig. 5C). Importantly, in association with reduced neuropathology, impaired spatial learning and memory of 5xFAD mice in the Barnes maze was partially restored by germline VGF overexpression (Fig. 5D).

**Fig. 5.**
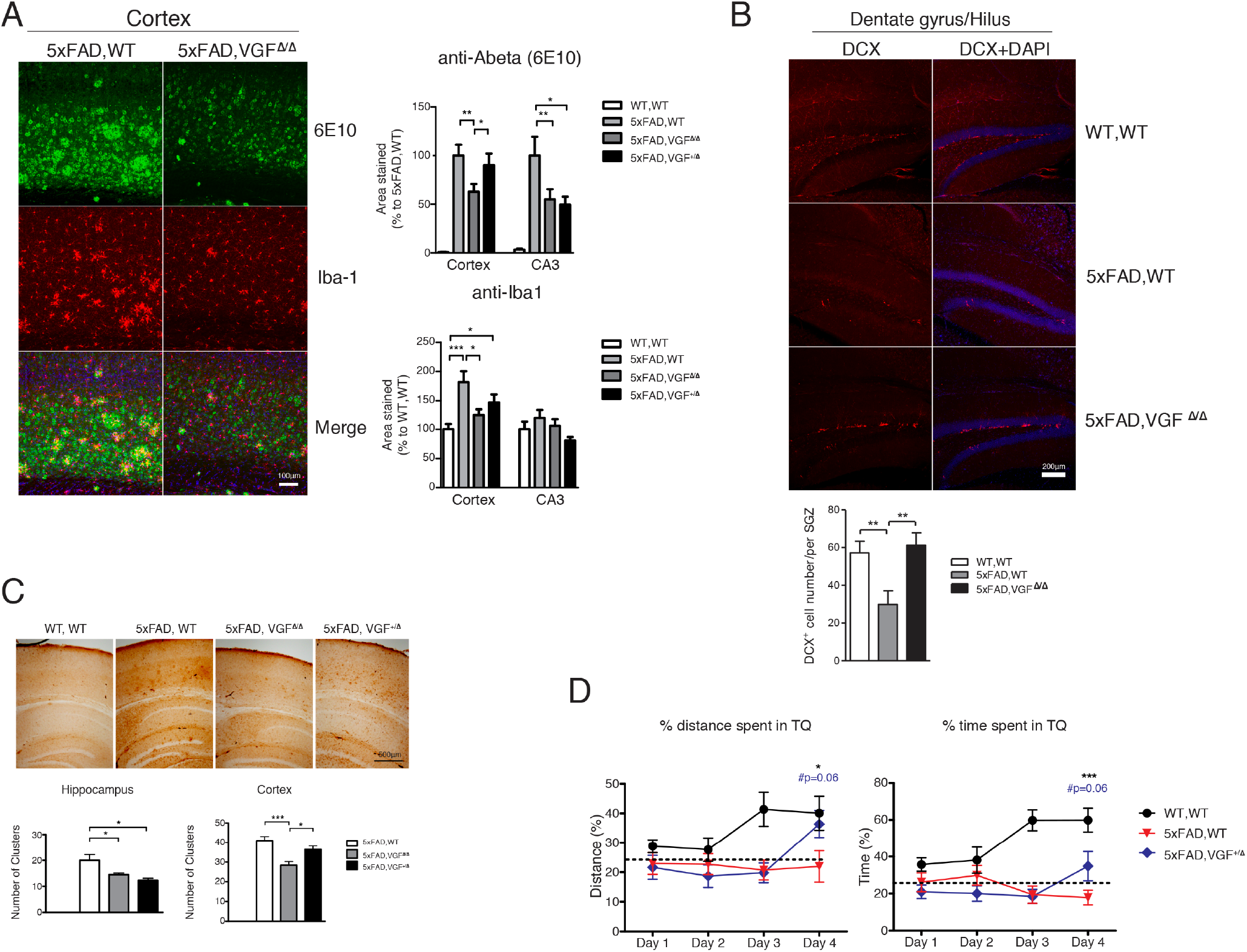
Characterization of AD pathophysiology in wildtype, 5xFAD, and 5xFAD mice overexpressing VGF. **A)** Immunohistochemical staining of Aβ amyloid plaques and microglial cells in the mouse cortex of 5xFAD mice overexpressing VGF in the germline. Left panel, green: Aβ (6E10), red: Iba-1, blue: DAPI; right panel, quantification of percent area of Aβ and Iba-1 staining. *N=6~9 male mice/per group.* **B)** Doublecortin staining (DCX) of the subgranular zone (SGZ) in the dentate/hilus area of 5xFAD brains. Upper panel, red: DCX, blue: DAPI; lower panel, average number of DCX-positive cells per subgranular zone. N=4 male mice/per group. **C)** Reduced staining of phosphor-Tau and dystrophic neurite clusters in 5xFAD brains with germline VGF overexpression. Upper panel: phosphor-Tau staining; lower panel: Quantification results of dystrophic neurite clusters in the hippocampus and cortical area. N=4~7 male mice/per group. Data of panel A, B, and C were analyzed by one-way ANOVA with Newman-Keuls post hoc analysis. *: p<0.05, **: p<0.01, ***: p<0.001. **D)** Barnes maze test. Mice were trained daily and wild type mice learned the target quarter (TQ) of the hiding zone by increased distance traveled in the TQ (left panel) and increased time spent in the TQ (right panel). 5xFAD mice showed impaired spatial learning on day 4, while germline VGF overexpression (5xFAD,VGF+/Δ) partially restored memory performance. N=12~14 mice (male+female)/per group. Data were analyzed by one-way ANOVA with Fisher’s LSD test. *: p<0.05, ***: p<0.001.

To examine whether overexpression of VGF in the adult 5xFAD brain also reduces neuropathology resulting from Aβ overexpression, adeno-associated virus (AAV)-VGF and AAV-GFP (control) were injected into the dorsal hippocampus (dHC) of adult 5xFAD mice at 2-3 months of age. We chose this brain region based on the following observations: i) previous studies indicated pro-cognitive effects of VGF peptide administration in the dHC in wild-type mice (*35*); ii) local VGF ablation in the mouse dHc resulted in memory deficits (*35*); iii) dHc’s proximity to the ectorhinal area, that, as described above, sustained the most damage by AD, with VGF identified in this region as significantly downregulated for multiple AD features. Animals were sacrificed for histological analysis at 7 months of age or at 10 months of age following behavioral testing. Immunohistological examination showed robust VGF overexpression transduced by AAV-VGF administration to the dHC of 5xFAD mice (Fig. 6A). Reduced levels of 6E10-immunoreactive plaques were found in the hippocampal dentate gyrus and nearby cortical regions (Fig. 6A). Similar to germline VGF overexpression, dHc AAV-VGF administration also restored neurogenesis in 5xFAD hippocampus to the level of wild type controls, and significantly reduced the number of dystrophic neurite clusters in the hippocampus (Fig. 6B, C). At 10 months of age, AAV-VGF-administered 5xFAD had significantly improved spatial learning and memory performance in the Barnes maze compared to those administered AAV-GFP, while VGF overexpression in non-transgenic wild type mice didn’t enhance memory, indicating a critical role for VGF in the pathological progression and behavioral impairment of the 5xFAD mouse model (Fig. 6D).

**Fig. 6.**
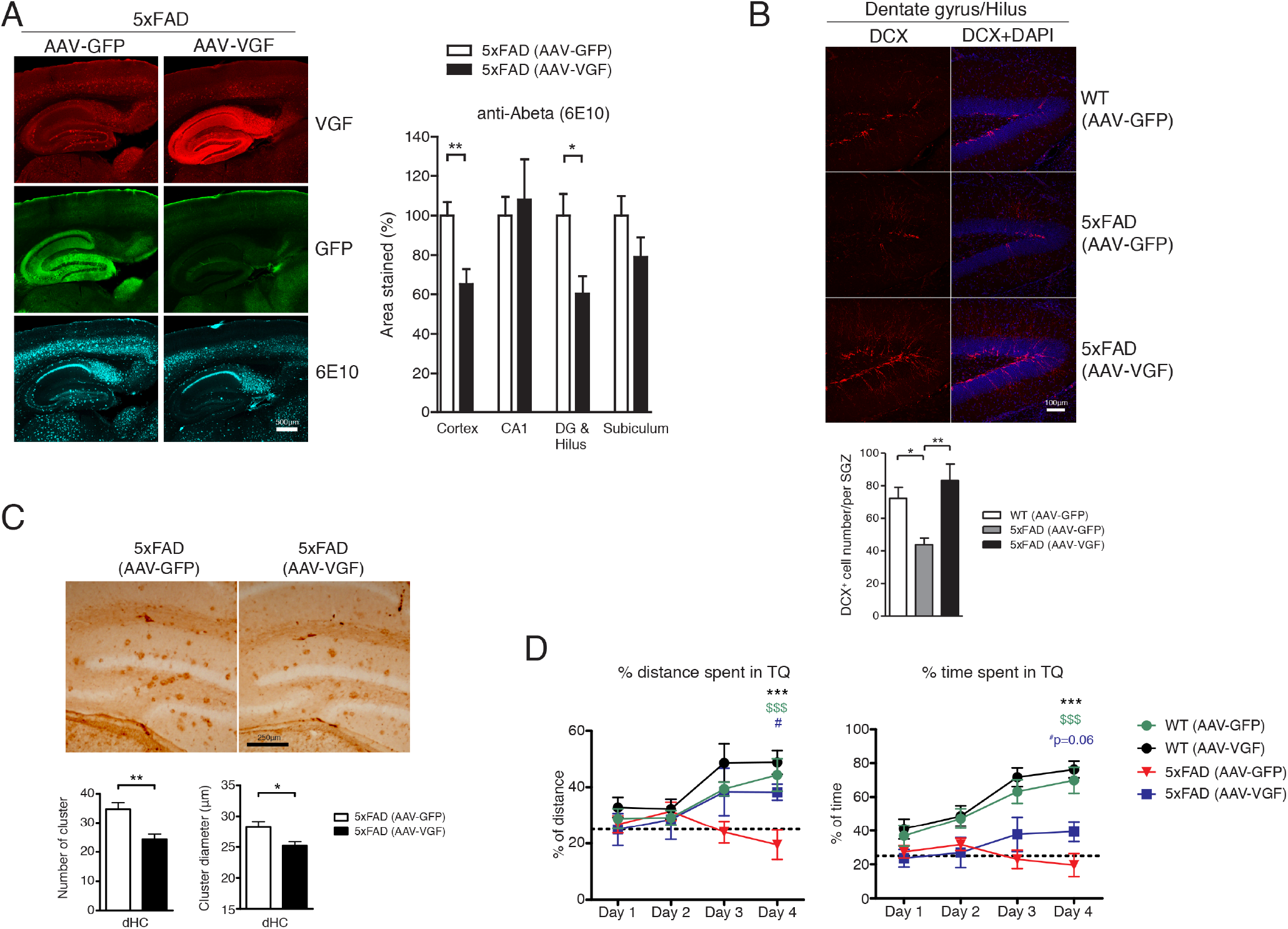
Characterization of AD pathophysiology in 5xFAD mice with and without AAV5-VGFdriven overexpression of VGF. **A)** Immunohistochemical staining of Aβ amyloid plaque*s* and VGF in the 5xFAD mouse brain 4 months after AAV5-VGF or AAV5-GFP infusion into dorsal hippocampus. Left panel, red: VGF, cyan: Aβ, green: GFP; right panel, quantification of percent area of Aβ amyloid plaque in different brain areas. N=4~5 male mice/per group. Data were analyzed by Student t-test. *: p<0.05, **: p<0.01. **B)** Doublecortin staining (DCX) in the dentate/hilus area. Upper panel, red: DCX, blue: DAPI; lower panel, average number of DCX-positive cells/per subgranular zone. N=4~5 male mice/per group. Data were analyzed by one-way ANOVA with Newman-Keuls post hoc analysis. *: p<0.05, **: p<0.01. **C)** Reduced staining of phosphor-Tau and reduction of dystrophic neurite cluster number and diameter in 5xFAD brains with AAV5-VGF overexpression. Upper panel, phosphor-Tau staining; lower panel, quantification results of dystrophic neurite cluster number and diameter in the dorsal hippocampus. N=4~5 male mice/per group. Data were analyzed by Student t-test. *: p<0.05, **: p<0.01. **D)** Barnes maze test. Mice were trained daily and on Day 4 wild type mice learned the target quarter (TQ) of the hiding zone, as revealed by increased *distance traveled* in the TQ (left panel), and increased time spent in the TQ (right panel). 5xFAD mice with AAV5-GFP showed impaired spatial learning on day 4, while in 5xFAD with AAV5-VGF overexpression, memory performance was significantly rescued. N=7~12 mice (male+female)/per group. Data were analyzed by one-way ANOVA with Fisher’s LSD test. *, #: p<0.05, ***, $$$: p<0.001.

VGF is a neuropeptide precursor that is processed into a number of bioactive peptides, including the C-terminal peptide TLQP-62 (named by the N-terminal 4 amino acids and length) (*92*). TLQP-62 has pro-cognitive and antidepressant efficacy and regulates neurogenesis, each of which is BDNF-dependent when the peptide is administered icv or directly to rodent hippocampus (*35*, *36*, *38*, *83*, *93*). Because BDNF is directly connected to VGF in our causal network, we investigated whether chronic 28-day icv administration of TLQP-62 to adult 3-4 month old 5xFAD reduced neuropathology at ~ 4.5 months of age. Significantly reduced levels of 6E10-immunoreactive plaques and Iba-1 immunostaining were found in hippocampal dentate gyrus and cortex of both TLQP-62-treated male and female 5xFAD (Fig. S7A, S7B), accompanied by significantly reduced numbers of Lamp1-immunoreactive dystrophic neurite clusters in hippocampus (Fig. S7C).

The pathophysiologic validation of VGF establishes this gene as one that can induce and protect against AD-related pathologies, as we predicted from our models. However, the pathophysiologic validation does not on its own confirm the molecular regulatory architecture of VGF defined by our network models. To validate VGF at the molecular network level, the brain gene expression signature induced in the mouse model by directly perturbing VGF (overexpressing this gene) can be compared to the genes predicted to change by the network in response to perturbations of VGF, as we have shown for different KD genes identified across a number of diseases (*5*, *15*, *18*, *24*, *25*, *27*). We sequenced RNA isolated from the prefrontal cortex of 89 mice with germline overexpression of VGF and corresponding controls, and from the hippocampus of 45 mice with AAV overexpression of VGF and corresponding controls. We found that genes downstream of VGF in the gene BN (predicted perturbation) were enriched for the AAV overexpression VGF DE signature (Data S12) at a threshold of FDR<0.05 (Fig. 7B, one-sided Fisher exact test OR=14.1, p-value=3.1e-6). We also found that while the germline overexpression VGF DE signature (Data S12) did not achieve significance at an FDR<0.05, DE genes at p-value<0.1 were also enriched downstream of VGF in the gene BN (Fig. S8, one-sided Fisher exact test OR=3.9, p-value=2.6e-3).

**Fig. 7.**
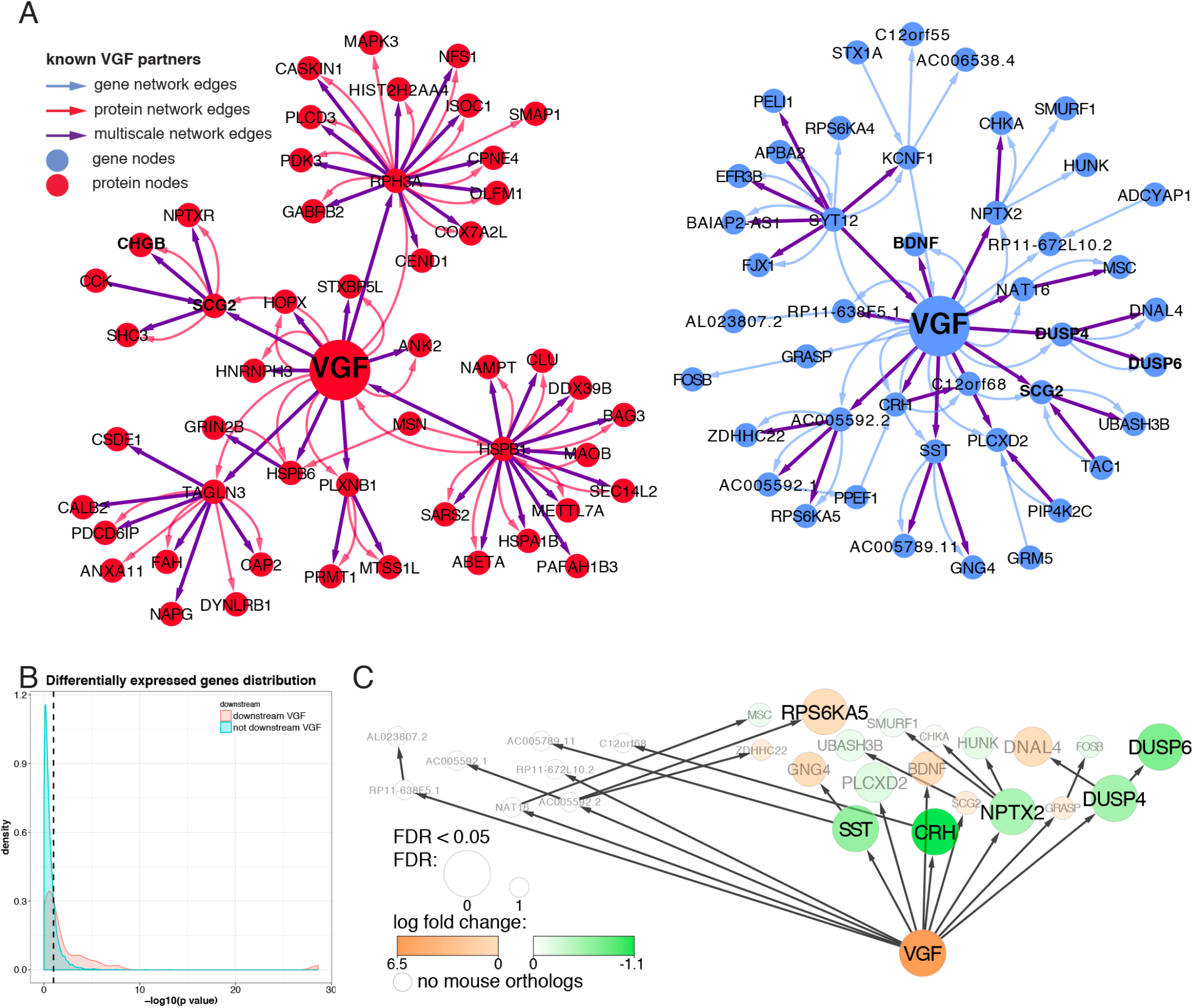
Molecular validation of VGF. **A)** Consensus subnetwork within a path length of 2 of VGF. The consensus subnetworks around VGF, 2 steps away from VGF, across all 3 networks are depicted. The blue and red nodes are genes and proteins, respectively. The blue edges originate from the gene only network, the red edges from the protein only network, and the purple edges from the multiscale network. VGF and its known partners are in bold in the plot. **B)** Density plot of the distribution of differential expression nominal p-values for genes downstream and not downstream (causally independent of the expression levels) of VGF in the gene-only network for mouse DE genes (5xFAD, AAV5-GFP versus 5xFAD, AAV5-VGF brains). The x axis is the −log10(p-value) for differential expression, and the y axis represents the densities at the different −log10(p-value). The red and blue curves are for genes downstream and not downstream of VGF in the network, respectively. **C)** Summary of DE results of VGF network genes in the 5xFAD, AAV5-GFP versus 5xFAD, AAV5-VGF brains overlaid on the VGF gene-only subnetwork. The nodes are colored by log fold change from green (negative) to orange (positive). The size of the node represents the DE FDR. Grey genes names are not significantly DE and white nodes have no orthologous genes in mice.

## Discussion

The primary aim of this project was to discover novel critical genes and pathways central to AD that could be potentially pursued as therapeutic targets. We applied a multiscale causal network modeling approach towards this end on the AMP-AD dataset in the anterior prefrontal cortex, which resulted in the identification of *VGF* as a novel key driver gene of AD. *VGF* was not only the most significantly downregulated gene in the protein and gene expression data in cases versus controls (best gene DE FDR = 5.0e-4, best protein DE FDR = 3.4e-15), but it was the only gene identified as a key driver across all three Bayesian causal networks we constructed. Further, *VGF* ranked as the top key driver gene not previously associated with AD, having the most explanatory power in distinguishing between AD cases and controls. We replicated VGF as a KD in an independent dataset as well as in 2 additional brain regions, demonstrated genetic support of its genomic locus associating with AD, and finally validated VGF *in vivo* at the physiologic and molecular levels as a driver of AD.

The biological coherence of the protein expression, compared to gene expression data, with respect to association with AD clinical features, was noteworthy, suggesting proteomic data may be a more informative measure for identifying important dysregulated pathways. Aβ, a hallmark of AD (*6*), was consistently the protein with the highest expression in cases relative to controls, whereas *VGF* was the most downregulated. DE proteins are annotated for energy metabolism and immune and nervous system related processes, all previously implicated in AD (*8*, *94*–*98*); our co-expression network analyses based in part on these DE protein studies, have further identified novel, potentially druggable targets within these pathways. As protein expression coverage increases and proteins with lower overall levels of expression are more reliably quantified, we can expect further improvements.

In a previous study of AD integrating transcriptomic and genomic data, TYROBP/DAP12 was identified as a major driver for the complement subnetwork during AD pathogenesis (*5*). In this instance, genotype and microarray gene expression measurements were processed to construct predictive network models (*5*). In the present study, we extended this approach by taking advantage of three layers of information, integrating genomic, transcriptomic and proteomic data to characterize the molecular association to AD and build predictive network models in order to causally infer relationships among these molecular traits and AD. The proteomic data not only adds a new dimension to this type of analysis, but the RNA sequencing data (compared to microarray data) provides a more comprehensive characterization of the functional units of the cell, thus allowing for a deeper modeling of the actual biological processes occurring in the cell and associated with disease. Interestingly, the immune-related protein module (green) containing VGF was significantly enriched for DE proteins across all clinical and neuropathological traits, emphasizing the importance of the immune system in AD and again highlighting the importance of the proteomic data in providing a more comprehensive characterization of AD.

The VGF gene we identified and validated is a nerve growth factor and brain-derived neurotrophic factor (BDNF) inducible gene (*99*) that is expressed in neurons in many different brain regions, and encodes a 615 amino-acid (617 in mouse) long precursor polypeptide (*100*) that is processed into several bioactive peptides that regulate neuronal activity and survival, neurogenesis, energy balance and lipolysis, and behavior (*35*, *36*, *100*–*105*). Moreover, induction of adult hippocampal neurogenesis in combination with elevation of BDNF levels, which occurs following exercise in rodents, has recently been shown to rescue cognitive deficits in 5xFAD mice (*106*). In this context it is important to note that VGF is robustly regulated in hippocampus by voluntary exercise (*107*) and by BDNF/TrkB signaling (*108*), and that in our studies, VGF overexpression rescued cognitive deficits and neurogenesis in 5xFAD mice (Fig. 5, 6). The VGF-derived peptide TLQP-62 (named by the four N-terminal amino acids and length), regulates neuronal activity, neural progenitor proliferation, memory formation, and depression-like behavior (*35*–*38*, *83*, *99*), via mechanisms that are largely dependent on BDNF/TrkB signaling (*35*, *36*, *109*) and was never showed to be causal to AD. In addition, TLQP-21, a sub-peptide of TLQP-62, activates the complement 3a receptor (C3aR1) (*103*, *104*); C3a activation of C3aR1 on microglia regulates amyloid uptake and microglial migration in primary microglia and/or mouse AD models (*110*–*112*). Lastly, the VGF1-617 proprotein, and secretogranin 2 (SCG2), identified in VGF gene and protein networks, are both granin genes and are linked through their functions in dense core vesicle (DCV) biogenesis and exocytosis, respectively (*113*).

A number of trait studies have found that VGF levels are reduced in the cerebrospinal fluid (CSF) of patients with AD (*40*, *42*–*44*, *46*, *114*, *115*). This is in agreement with our findings, where VGF is the gene and protein product with the lowest expression in cases relative to controls. Longitudinal measurement in patients and controls previously showed that VGF levels decrease with time as the disease progresses (*42*). Interestingly, reduced VGF levels were detected prospectively in CSF from patients with mild cognitive impairment, selectively in those who develop AD (*44*). VGF has also been proposed as a biomarker of AD and its progression (*42*). Although CSF levels of VGF, a neuronal and neurosecretory protein, might be anticipated to decrease coincident with neuronal loss as AD progresses, it is important to note that in two biomarker studies, the CSF levels of a number of related neurosecretory and synaptic proteins, including chromogranin A, secretogranin II, 7B2, proSAAS, clusterin, neurexins 1, 2 and 3, and neuropentraxin 1 were either increased or unchanged in patients with AD compared to controls, while VGF levels consistently were reduced in AD patient CSF (*41*, *44*). Our systematic analysis of changes in gene expression and proteomic profiles in disease-free and AD brains identified VGF as not only strongly associated with AD, but as a key driver gene of our AD-associated network, a finding that is further supported by our direct perturbation of VGF in the 5xFAD mouse model. Notably, many genes that are one or two steps downstream of VGF in our network are CREB-responsive genes (FosB, Nptx2, Tac1, Scg2, SST, DUSP4, BDNF, CRH, PLCXD2, NPTX2), as is VGF, with one or more CREB-binding sites localized in the promoter region or gene expression induced by overexpression of constitutively active CREB (*116*). Previous studies have also shown that VGF-derived peptide TLQP-62 activates the CREB signaling pathway in both acutely treated hippocampal slices or chronically infused rodent hippocampus, supporting VGF’s role as a key driver of these CREB-responsive network genes (*35*, *109*).

The majority of VGF network genes regulate neuronal activity, synaptic plasticity, and cognitive function. Previous reports indicate that levels of proteins encoded by the SST, NPTX2, SCG2, BDNF, and DUSP6 genes are reduced in the AD brain (*117*–*121*). BDNF, CRH and SST modulate neuronal activity and synaptic function, while BDNF and CRH also show neuroprotective effect against Aβ insults (*122*, *123*). DUSP4 and DUSP6 belong to the dual specificity phosphatase family (DUSPs). DUSP4 knockout mice have spatial reference and working memory deficits (*124*), while DUSP6 is expressed in microglia and is regulated by BDNF gene ablation in PFC (*125*, *126*). Both SCG2 and VGF are components of DCVs and their levels are critical to the biogenesis and regulated secretion of DCVs, therefore, their imbalance in the AD brain may cause dysregulation of neurotrophin, neuropeptide and/or catecholamine secretion and function (*113*). Genes in the VGF network that have particular relevance to the regulation of neural activity include GRASP, a scaffold protein that is involved in intracellular mGluR trafficking (*127*), and NPTX2 (NARP), reduction of which results in GluA4 reduction and further disruption of hippocampal gamma oscillation in the AD/NPTX2 KO mouse model (*118*).

The VGF-driven protein only network identified in our study reinforces the important role that homeostatic levels of VGF play in the regulation of neuronal integrity and plasticity. Among the proteins directly connected to and downstream of VGF in our network, critical roles for ANK2, PLXNB1, TAGLN3 and HSPB6 have been identified in the regulation of dendritic and axonal morphology as well as cytoskeletal reorganization (*128*–*133*). Other proteins directly connected downstream of VGF were SCG2, required for neuronal differentiation and maturation (*134*), HOPX, modulates hippocampal neurogenesis (*135*), and STXBP5L and RPH3A, which are involved in the trafficking and release of neuronal synaptic or dense core vesicles (*136*, *137*). Relevant to our analysis, previous protein crosslinking studies have identified a VGF interaction with amyloid precursor-like protein 1 (APLP1) (*138*). Whether this interaction mechanistically impacts VGF or b-amyloid function in AD brain is unknown, but interestingly, preliminary immunohistochemical analysis has visualized VGF but not NPY immunoreactivity associated with amyloid plaques in 5xFAD hippocampus (Lin W.J., Hariharan S., unpublished data).

While the determination of the precise mechanism(s) of action of VGF in AD requires additional study, constructing and validating models of AD that can serve as a more integrated and comprehensive repository of the regulatory frameworks of AD, provides a more informative and accessible path for others to leverage extensive sets of data from which they can validate links between known disease targets, generate hypotheses around novel targets, and derive mechanistic insights that further our understanding of AD. Differences between specific molecular mechanisms and subsets of disease are of great interest for further exploration of AD networks, however the focus of this work was on constructing a predictive model of AD and validating the top master regulator identified of the networks. Indeed, the data presented here are consistent with causal roles for TLQP-62 and the VGF proprotein in Alzheimer’s disease pathogenesis and progression, but do not rule out contributions of other VGF-derived peptides including TLQP-21, an activator of the C3aR1 complement receptor (*103*, *104*), that will require additional investigation.

## Materials and Methods

### Data description

All MSBB discovery and replication datasets were previously described in Wang et al. (*47*). These consist of gene and protein expression and WES for a cohort of individuals across the entire spectrum of AD in the Mount Sinai Brain Bank. RNA-seq was performed for 1096 samples from 315 individuals across 4 brain regions, and MS/MS for 266 samples from 266 individuals in one of the brain regions to measure protein expression. Whole exome was sequenced for 309 individuals. The RNA for all samples was treated with Ribo-Zero to remove rRNA and keep other transcripts (*139*). The 4 brain regions assessed are the anterior prefrontal cortex (BM10), the superior temporal gyrus (BM22), the perirhinal cortex (BM36) and the pars opercularis (BM44). Protein expression was restricted to BM10. The disease was categorized in 6 different ways, each representing different aspects of AD: clinical dementia rating (CDR), clinical neuropathology (Path Dx), CERAD neuropath criteria (CERJ), neuropathology category (NP-1), mean neocortical plaque density (PlaqueMean, number of plaques per mm^2^), and Braak score (bbscore) (*140*–*145*). The religious order study and memory aging project (*48*, *49*) (ROSMAP) validation set consists of gene expression from the dorsolateral prefrontal cortex of 724 subjects and whole genome sequencing (WGS) data from 1200 subjects. The ROSMAP RNA-seq count matrix and associated quality measurements were downloaded from https://www.synapse.org/#!Synapse:syn9702085, where their generation is described. The ROSMAP WGS data variant call format (VCF) file was downloaded from https://www.synapse.org/#!Synapse:syn10901595, where its generation and quality control are described. All data are available at https://www.synapse.org/#!Synapse:syn2580853/wiki/409853

### RNA-seq processing

In order to ensure a reliable set of samples and genes for all analyses in the MSBB datasets, we performed quality control processing and filtering for lowly expressed genes on the whole dataset across all 4 brain regions. Starting with the raw RNA-seq reads, we aligned (STAR) to GRCh37 and counted the reads mapping to each gene (featureCounts) as well as created QC matrix and called variants (GATK) on the RNA-seq with the RAPiD pipeline (*53*, *146*–*148*). For RNA-seq samples sequenced multiple times, we started by selecting the fastq file (raw reads) based on which had the largest number of mapped reads and less than 5% rRNA mapped reads. we then ran STAR alignment and featureCounts to generate the raw count matrix. we called variants on the RNA-seq data using GATK. Variants were also called for the WES using GATK. Using RNA expression and variants from the WES data, we imputed sex information for each sample. Comparing the heterozygous variants from the RNA-seq data to the variants in the WES data, enabled us to assign each RNA-seq sample to its corresponding DNA sequence. Using these multiple layers of information, we corrected, when necessary, mislabeling. For RNA-seq samples with documented matching WES, if the discordance rate between said sample and its best corresponding exome sequence was more than 10%, they were removed from further analyses. This left 958 RNA-seq samples in the MSBB dataset. In the ROSMAP gene expression data, we found one sample were gender was mislabeled and removed it from further analyses.

In the MSBB data, to filter out low expressed genes, we removed all genes that did not have at least 1 count per million (cpm) in at least 10% of the samples. we normalized the raw counts using the voom function from the limma R package (*149*–*151*). After exploration of the main drivers of variance using principal component (PC) analyses and using linear mixed models, we adjusted the normalized counts for batch effects using linear mixed models (variancePartition) (*54*). The corrected residuals were further adjusted with the lmFit function of the limma package for post mortem interval (PMI), race, sex, RNA Integrity Number (RIN) and Exonic Mapping Rate (*151*). Outlier samples, further than 3 standard deviations from the centroid of PC1 and PC2, were removed from downstream analyses. From the remaining samples, samples with RIN<4 were removed from further analyses. The raw counts of the 886 samples remaining were then subjected to the exact same protocol, to get normalized and adjusted gene expression for 24865 genes.

In the ROSMAP data, we followed a similar protocol removing all genes that did not have at least 1 cpm in at least 10% of the samples, normalized using the voom function and after exploration of the main drivers of variance, adjusted the normalized counts for Batch, sex, race, PMI, RIN, median 5 prime to 3 prime bias, strand balance, and percent of intronic bases using the lmFit function. The output was a matrix of normalized and adjusted counts of 19452 genes for 633 samples.

### Protein expression processing and correction for other covariates

The protein expression data was taken through similar procedures to ensure that there would be no technical variance in the way of true biological signal. After correction for Batch on the protein expression data for 266 samples, we further adjusted for PMI, race and sex using the lmFit function of the limma package (*151*). The remaining 2692 protein expression residuals were used for downstream analyses.

### Differential Expression analyses

The DE analyses were performed for both gene and protein expression using the limma package after the adjustment for covariates described earlier (*151*). In order to capture all aspects of the disease, the DE was performed for each AD trait. In addition, to capture signal corresponding to the entire spectrum of AD, DE analysis was performed in 2 ways: controls against any sample that had any level of cognitive impairment (and in the case of PlaqueMean using its quantitative level as a response), and definite controls against definite AD as defined by each trait (Table S1).

### QTL analyses

All QTL analyses were run using the fastQTL package (*154*). Starting from the variants from the WES (MSBB) or WGS (ROSMAP), using plink2, we removed markers with more than 5% missing rate, less than 1% major allele frequency and Hardy-Weinberg p-value lower than 10^−6^ (*155*, *156*). Following standard practice, only European individual were used to find QTLs (*157*). Non-European samples were identified through PCA analyses using smartPCA and mapping in PC space to the 1000 Genomes Project consortium (*158*, *159*). VCF-liftover was used to lift over the ROSMAP WGS from hg19 to hg38 (*160*). The residuals described above were used for QTL analyses for both gene and protein expression after further correction for PEER surrogate (latent) variables variables (SVs) (*161*) as follows: (i) BM10 gene expression: 19 SVs; (ii) BM10 protein expression: 9 SVs; (iii) BM22 gene expression: 20 SVs; (iv) BM36 gene expression: 17 SVs; (v) BM44 gene expression: 17 SVs; (vi) ROSMAP gene expression: 25 SVs. We also included in the model the first 5 PCs of the genotype data to remove further population specific structures. The analyses looked for cis-eQTL as defined 1 Mb of the transcription start site of each gene and protein corresponding gene. FDR were computed following Benjamini-Hochberg (*162*). The causal inference testing was performed with the R package citpp (https://bitbucket.org/account/signin/?next=/multiscale/citpp).

### Co-expression analyses

Two WGCNA co-expression networks were built on the adjusted data, one for the gene expression and one for the protein expression, using the coexpp R package (*163*) (Michael Linderman and Bin Zhang (2011). https://bitbucket.org/multiscale/coexpp). To identify modules of interest in the context of AD, we projected the union of all DE genes or proteins on the corresponding co-expression network. We calculated enrichment statistics using Fisher’s Exact Test, and corrected for multi-testing following Benjamini-Hochberg.

### Seeding gene list construction

Making the assumption that DE genes are important for AD, and that therefore these genes need to be included in the model, we started by adding the union of all DE genes to the seeding gene list. To include other important genes that co-vary with these DE genes, but that may not reach significance in the DE analyses, we included all gene in co-expression modules enriched for DE genes. Finally, for the discovery gene expression set only (BM10), to maximize the chances to not miss important genes, we added to the gene list of DE genes and modules of interest other genes known to be connected to our current gene list in the literature using PEXA (*74*). In order to build the extended network, PEXA used KEGG pathways, and to trim it, used a PPI network from CPDB (*164*, *165*). We used the outputted discovery seeding gene list of 5714 genes for which we have gene expression for the purpose of Bayesian causal network construction. For the replication sets, the seeding gene lists were comprised of only the DE genes and the coexpression modules enriched for the DE genes, and consisted of respectively 10585 genes for BM22, 16578 genes for BM36, 8086 genes for BM44 and 9682 genes for ROSMAP. For BM44, due to the small number of DE genes and to increase the number of genes in the AD DE gene set with genes related to DE genes and find co-expression module enrichments, we added the top 10 correlated genes to each of the DE genes before module enrichments.

### Bayesian Causal Networks

BNs were built using RIMBANET (*32*–*34*, *51*) using gene expression, protein expression and both gene and protein expression (multiscale) for the discovery datasets, and using gene expression for the replication datasets. In each case, QTLs were used as priors (eQTLs for gene networks, pQTLs for protein network and both for multiscale network). To reduce the search space and increase the likelihood to reach a global maximum of the fit of the network, we reduced the gene space from entire expressed transcriptome (24865 genes) to the seeding gene list described earlier for the gene only and the multiscale networks. Because there was protein expression for only 2692 proteins, we included all the protein in both the protein and the multiscale networks. Because of the central dogma of biology and the results of the CIT analysis, we included strong weak edge priors (increasing the likelihood for that edge to be searched) to the multiscale network from the parent gene to its corresponding protein product. Representation of networks and subnetworks was achieved using the Cytoscape software version 3.5.1 (*166*).

### Key Driver Analyses

For both the discovery and the replication datasets, to do Key Driver Analysis, we used the R package KDA (*75*) (KDA R package version 0.1, available at http://research.mssm.edu/multiscalenetwork/Resources.html). This package defines a background sub-network by looking for a neighborhood K-step away from each node in the target gene list in the network. Stemming from each node in this sub-network, it assesses the enrichment in its k-step (k varies from 1 to K) downstream neighborhood for the target gene list. In this analysis, we used K=6. KD analyses were performed by projecting multiple seeding target lists of interest on the networks: The DE lists of the corresponding omics for each disease trait to find KDs of the diseases. In the discovery dataset, KD were then prioritized by first how many networks they appear in (replication) and then how many times they appear across networks (importance). In the replication datasets, we looked for presence of VGF as a KD of the networks.

### Ranking of KDs using Machine Learning

For each classifier we performed a random split of the data, stratified by class, into 75% training set and 25% validation set (*167*). The training set was subjected to SMOTE (*168*, *169*) to resolve any class imbalance for training the random forest (RF) classifier (python sklearn package (*170*, *171*)). Classifier performance was evaluated against the validation set and quantified using area under the curve (AUC) of the receiver operating characteristic (ROC) curve (*172*). RF randomly sub-sets the features into decision trees, selecting a feature from each subset that best separates the data into classes (*170*). Therefore, the choice of a feature to be included in the forest is an indication of the performance and stability of that feature. Features were ranked by importance, as based on information gain score (*173*). This process was performed 500 times to estimate the distribution of feature information gain across classifiers. Features were then organized into a meta-rank by a weighted z-score method across the 500 iterations per classifier (*174*, *175*). There, a z-score was established from the features rank per iteration of the information gain and weighted by a factor accounting for the stability of features and the performance of the classifier.

The weight is the product of two components: (i) the ratio of the number of iterations each feature appeared in to the mean number of all features iteration appearances, both across the 500 forests; (ii) the absolute value of the ROC AUC score of each classifier centered at 0, minimizing the impact of random classifiers.

The classifiers were run independently, after normalization and adjustment for covariates, on each scale of expression data: the gene expression data, the protein expression data and the gene and protein expression data together. In each case, the 11 AD traits were used as classes to train and test the classifiers and a meta-rank for each feature across the 11 traits was computed using the weighted z-score approach described earlier across all 5,500 classifiers. To further prioritize the networks KD, all KDs were ordered according to the meta-rank of features across traits for their corresponding scale of data.

### Manhattan Plot of GWAS results

Summary statistics for the largest AD GWAS to date (17,008 and 37,154 controls, I-GAP GWAS) results (*10*) around the VGF gene (+/‐ 125 KB) were plotted using LocusZoom (*176*).

### Other statistical analyses

R version 3.3.1 was used for statistical analyses unless specified otherwise (*177*). GO annotations enrichment was tested with, the R packages goseq (*178*), topGO (Alexa A and Rahnenfuhrer J (2010). topGO: topGO: Enrichment analysis for Gene Ontology. R package version 2.18.0.) and org.Hs.eg.db (Carlson M. org.Hs.eg.db: Genome wide annotation for Human. R package version 3.2.3.). To test MSigDB pathway enrichment, the R packages HTSanalyzeR (*179*), GSEABase (Morgan M, Falcon S and Gentleman R. GSEABase: Gene set enrichment data structures and methods. R package version 1.32.0.), and gage (*180*) were used. Figures where generated using the R packages ggplot2 (*181*), scales (Hadley Wickham (2012). scales: Scale functions for graphics. R package version 0.2.3. http://CRAN.Rproject.org/package=scales), reshape2 (*182*) (http://www.jstatsoft.org/v21/i12/.) and grid (*183*). UpsetR plots were generated with the UpSetR R package (*184*). Heatmaps were produced with the function heatmap.2 of the R package gplots (*185*). Venn diagram were dawn using the VennDiagram R package (*186*). Circos (circular) plot of DE enrichments in modules were plotted using the NetWeaver R package (*153*, *187*). Canonical Correlation analyses were performed with the canCorPairs function of the variancePartition R package (*54*).

### Animal models and stereotaxic surgery

The generation of 5xFAD mice was described previously (*81*). These transgenic mice overexpress both human APP (695) harboring the Swedish (K670N, M671L), Florida (I716V) and London (V717I) familial AD (FAD) mutations and human Presenilin1 (PS1) harboring the two FAD mutations M146L and L286V. Expression of both *trans*-genes is regulated by neuronal-specific elements of the mouse *Thy1* promoter. The 5xFAD strain (B6/SJL genetic background) was maintained by crossing hemizygous transgenic mice with B6/SJL F1 breeders. The floxed VGF mouse line was generated as recently described (*82*). Homozygous floxed VGF mice that overexpress VGF mRNA and protein by virtue of the placement of the pgk-neo cassette in the 3’ UTR region of the *Vgf* gene. This leads to premature mRNA termination and polyadenylation utilizing a cryptic poly-A addition site in the inverted pgk-neo cassette, truncating part of the 3’UTR sequence, and resulting in increased CNS expression of VGF (Fig. S5). All mouse studies were conducted in accordance with the U.S. National Institutes of Health Guidelines for the Care and Use of Experimental Animals, using protocols approved by the Institutional Animal Care and Use Committee of the Icahn School of Medicine at Mount Sinai.

Mice at 2 – 3 months of age were anesthetized with a mixture of ketamine (100mg/kg) and xylazine (10mg/kg). Thirty-three gauge syringe needles (Hamilton, Reno, Nevada) were used to bilaterally infuse 1.0 μl of AAV virus into mouse dorsal hippocampus (dHc) (AP = −2.0, ML = ±1.5, and DV = −2.0 from Bregma (mm)) at a rate of 0.2 μl per min and the needle remained in place for 5 min before removal to prevent backflow. AAV5-GFP and AAV5-VGF (mouse VGF cDNA) were prepared by the Vector Core at the University of North Carolina at Chapel Hill. AAV-injected mice were used at 7~8-month old for immunohistochemical analysis or at 10-month old for behavioral analysis. Additional mice at 3 months of age were anesthetized with ketamine/xylazine and a cannula was implanted in the lateral ventricle [AP=−0.1, ML=±1.0 and DV: −3.0 from bregma (mm)] (*188*). TLQP-62 (2.5 mg/ml) dissolved in aCSF or aCSF alone was delivered icv by microosmotic pump (Alzet delivering 0.25 μl/h or 15 μg/day) for 28 days. Mice were used for immunohistochemical analysis at 4.5 months of age.

### Immunohistochemical and biochemical analysis

Immunohistochemical and biochemical characterization were performed as previously described (*3*, *189*–*192*). For biochemical analysis, hemibrains were processed via differential detergent solubilization to produce TBS-soluble, Triton-X-soluble, and formic-acid soluble Aβ fractions. For analysis of native oligomeric Aβ peptides, 2 μl protein samples from the TBS-soluble fraction were spotted onto activated/pre-wetted PVDF membrane (0.22 μm; Millipore, Billerica, MA). Membranes were incubated with rabbit pAb A11 (anti-prefibrillar oligomers, 0.5 μg/ml), rabbit pAb OC (anti-fibrillar oligomers and fibrils; 0.25 μg/ml), and mouse mAb Nu-4(anti oligomers; 1 μg/ml) (*191*, *192*). Normalization to total APP/Aβ signal was achieved by detection of human APP transgene protein with the mouse pAb 6E10 antibody (1:1000; Covance, Princeton, NJ). To quantify total Aβ levels, human/rat Aβ 1–40/1–42 ELISA kits (Wako, Richmond, VA) were used according to the manufacturer’s instructions. For immunohistochemistry, 50 μm thick sagittal sections were incubated with the following antibodies: rabbit anti-Iba1 (1:500; Wako, Richmond, VA), mouse anti-6E10 (1:1000; Covance, Princeton, NJ), rabbit anti-doublecortin (1:500, Cell signaling Technology, MA). Sections were then incubated with appropriate secondary antibodies: anti-mouse Alexa Fluor 488 (1:500; Invitrogen, Carlsbad, CA), anti-rabbit Alexa Fluor 594 (1:500; Invitrogen, Carlsbad, CA). For non-fluorescent immunostaining, endogenous peroxidase was quenched with PBS containing 3% H_2_O_2_, followed by amplification using the ABC system (VECTASTAIN Elite ABC HRP Kit, Vector Laboratories, Burlingame, CA). Horseradish peroxidase conjugate and 3,3′-diaminobenzidine were then used according to the manufacturer’s manual (Vector DAB, Vector Laboratories, Burlingame, CA). ThioflavinS (Sigma-Aldrich, T1892, 1% w/v stock solution) was used for labeling amyloid deposits. For immunoblotting, membranes were incubated with either anti-VGF C-terminal (1:1000; rabbit polyclonal), anti-6E10 antibody (1:1000; Covance, Princeton, NJ), anti-actin (1:1000; Sigma-Aldrich) antibodies. The membranes were washed, incubated with a secondary horseradish peroxidase-labeled donkey anti-rabbit or donkey anti-mouse antibody (1/6000; GE Healthcare) for 1 h, washed again, and incubated with ECL detection reagents (Millipore). Densitometric analysis was performed using ImageJ software.

### RNA extraction and qPCR analysis

RNA from mouse tissue specimens, obtained by dissection (prefrontal cortex), was extracted using miRNeasy Mini Kit (Qiagen) according to the manufacturer’s protocol, and 0.25 μg was reverse transcribed using iScript reverse transcription supermix for RT-qPCR kit (Bio-Rad, Hercules, CA). One nanogram of first-strand cDNA was subjected to PCR amplification using a SYBR green real-time reverse transcription PCR (qPCR) master mix (PerfeCTa SYBR Green FastMix, Quanta Biosciences). ΔΔCt method was used to quantify relative gene expression and normalized to glyceraldehyde 3-phosphate dehydrogenase (Gapdh).

### Behavioral test and analysis

The Barnes Maze test was performed using a standard apparatus (*193*, *194*). 10-month-old mice were transported from their cage to the center of the platform via a closed starting chamber where they remained for 10 s prior to exploring the maze for 3 min. Mice failing to enter the escape box within 3 min were guided to the escape box by the experimenter, and the latency was recorded as 180 s. Mice were allowed to remain in the escape box for 1 min before the next trial. Two trials per day during 4 consecutive days were performed. The platform and the escape box were wiped with 70% ethanol after each trial to eliminate the use of olfactory cues to locate the target hole. All trials were recorded by video camera and analyzed with ANY-maze video tracking software (Stoelting Co, Wood Dale, USA).

## Authors Contributions

NDB, WJL, MW, BZ, SRS and EES conceived, designed and managed the study with advisory input from ME. NDB performed all statistical and computational analyses under the direction of EES, with advisory input from ATC. WJL performed all biological experiments under the direction of SRS, with experimental support from CJ, SH, MA, JVHM, BS, SG and ME. PW and WM normalized and input protein expression. ATC, YCW, GMB, PC, AWC, EEK, HZ, and ZT provided critical technical and computational support. AIL, NTS, EBD, DD and JJL performed protein sequencing, quantification and advised on protein expression processing. VH and PK collected the brain tissue used in the study. NDB, WJL, SRS and EES wrote the manuscript, with input from MW, BZ and ME. All authors critically reviewed and edited the manuscript. All authors contributed significantly to the work presented in this paper.

## Data and material availability

all data and materials are available either in the main text and supplementary materials, or in www.synapse.org/ampad.

---

**Fig. S1.**
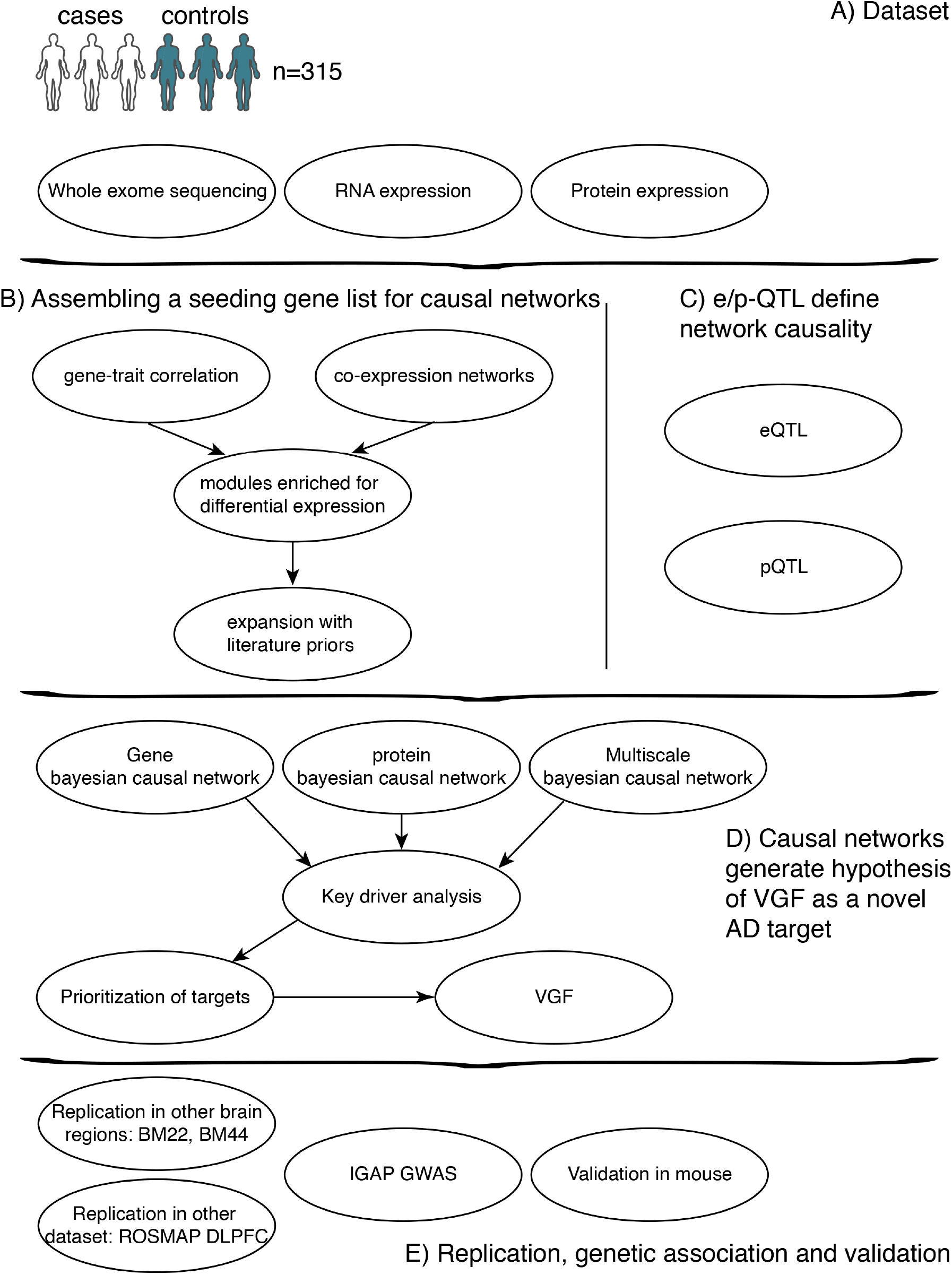
Simplified Pipeline Overview. Simplified description of data and analyses workflows performed to identify and validate VGF as a target of AD.

**Fig. S2.**
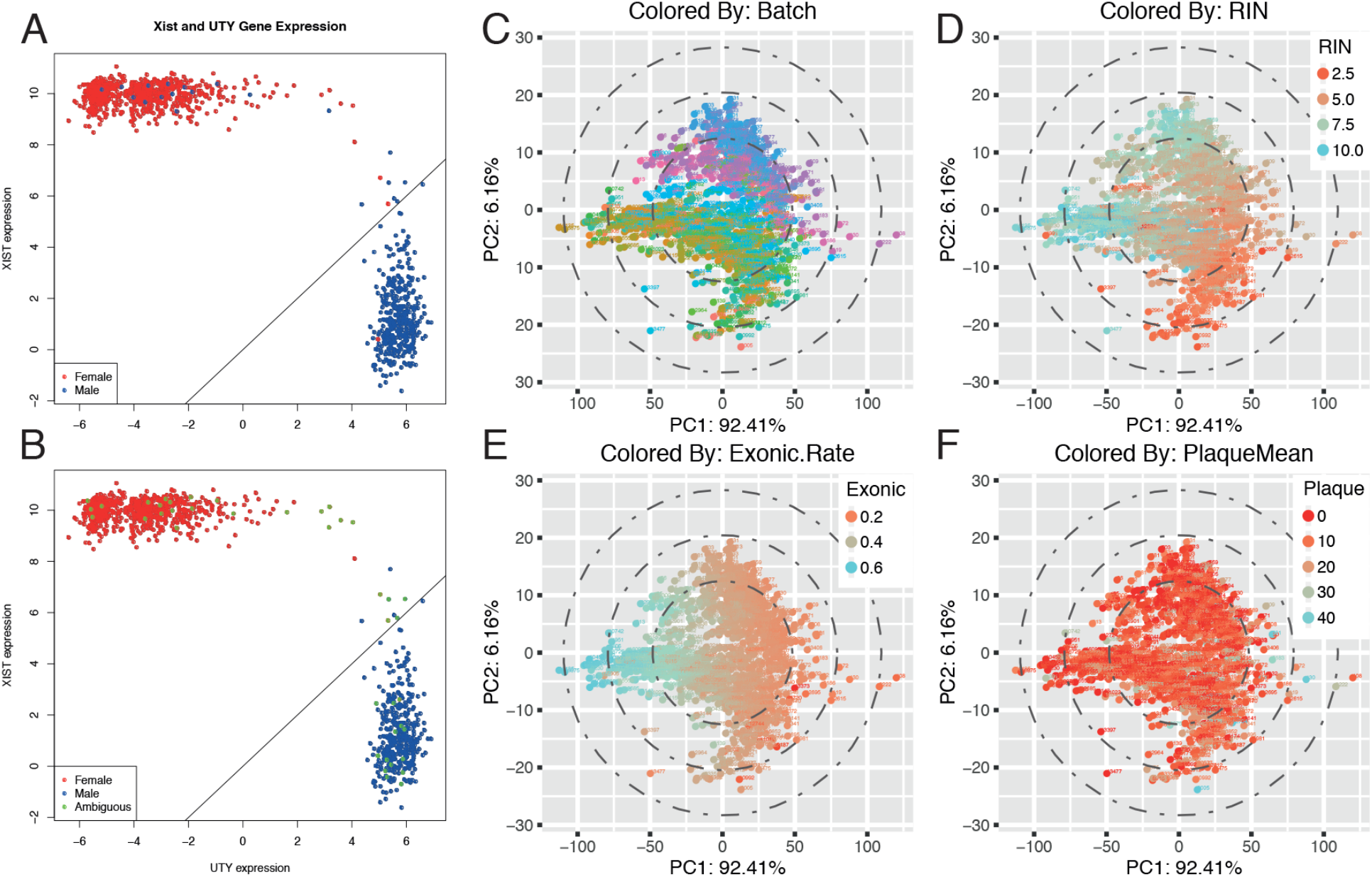
Data quality control. **A and B)** Imputed RNA-seq sex colored by sex clinical information: Normalized gene expression for XIST (female specific gene, y axis) and UTY (ubiquitously expressed Y-chromosome gene, male specific, x axis). **(A)** Obvious sex mislabeling is present in the dataset. **(B)** After fixing the mislabeling, ambiguous samples (removed from further analyses) are shown in green. **C, D, E and F)** Principal component analyses of important covariates: panels of this figure represent the same samples (one sample per point). The x axis is PC1 and explains 92.41% of the variance in the expression data. The y axis is PC2 and explains 6.16% of the variance in the expression data. The samples are colored by different QC or clinical information associated to them.

**Fig. S3.**
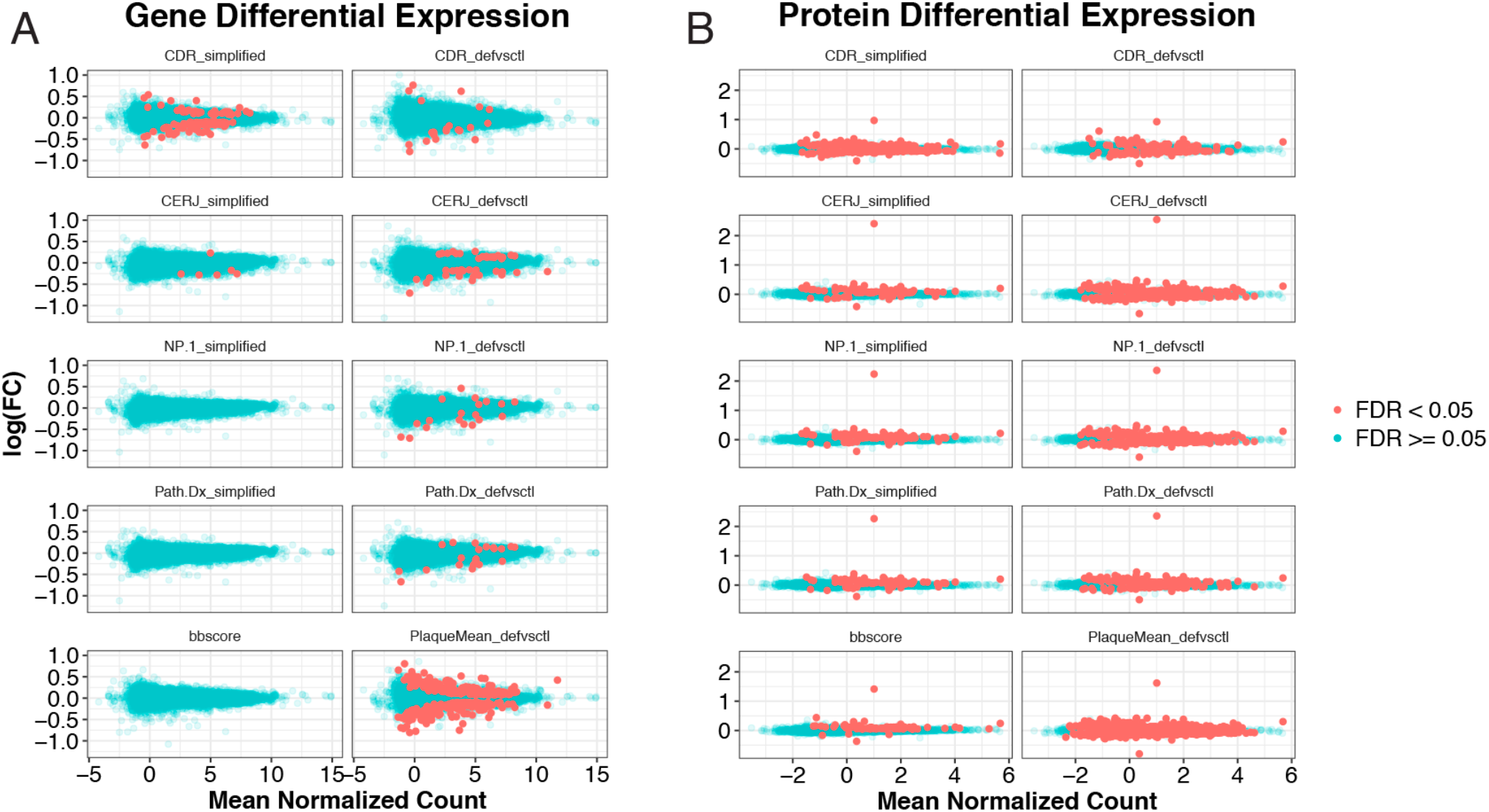
Other AD traits gene DE. **A and B)** Gene **(A)** and protein **(B)** differential expression: The x axis of this plot is the mean normalized count for each gene or protein, and the y axis the log(FC). In blue are the non-significantly DE genes or proteins and in red the significant ones. Each box corresponds to a trait

**Fig. S4.**
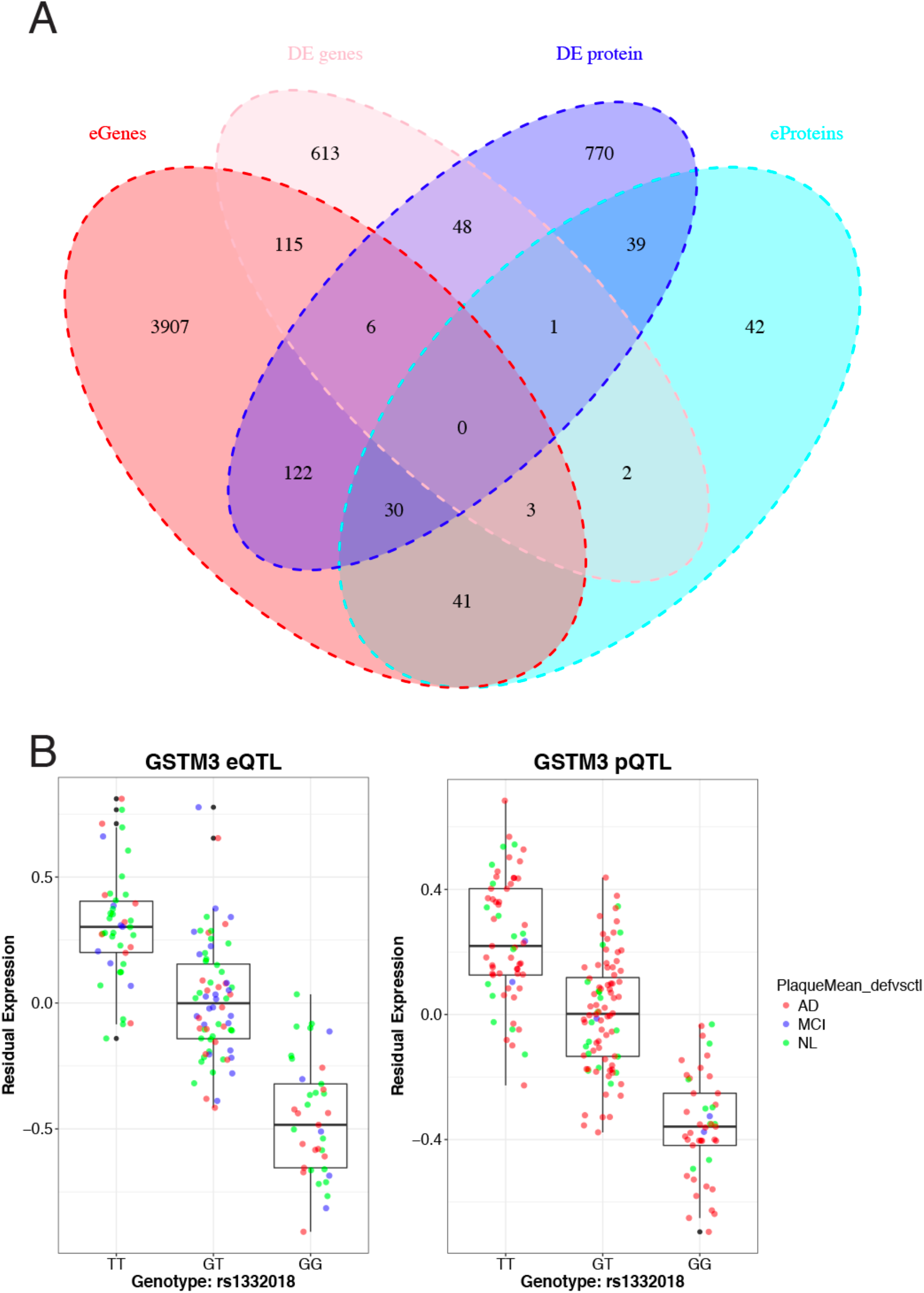
QTL analyses. **A)** Venn diagram of QTL overlap with expanded DE signatures. **B)** Boxplots of QTL effects: GSTM3, gene that shares an eQTL and a pQTL at the same SNP position.

**Fig. S5.**
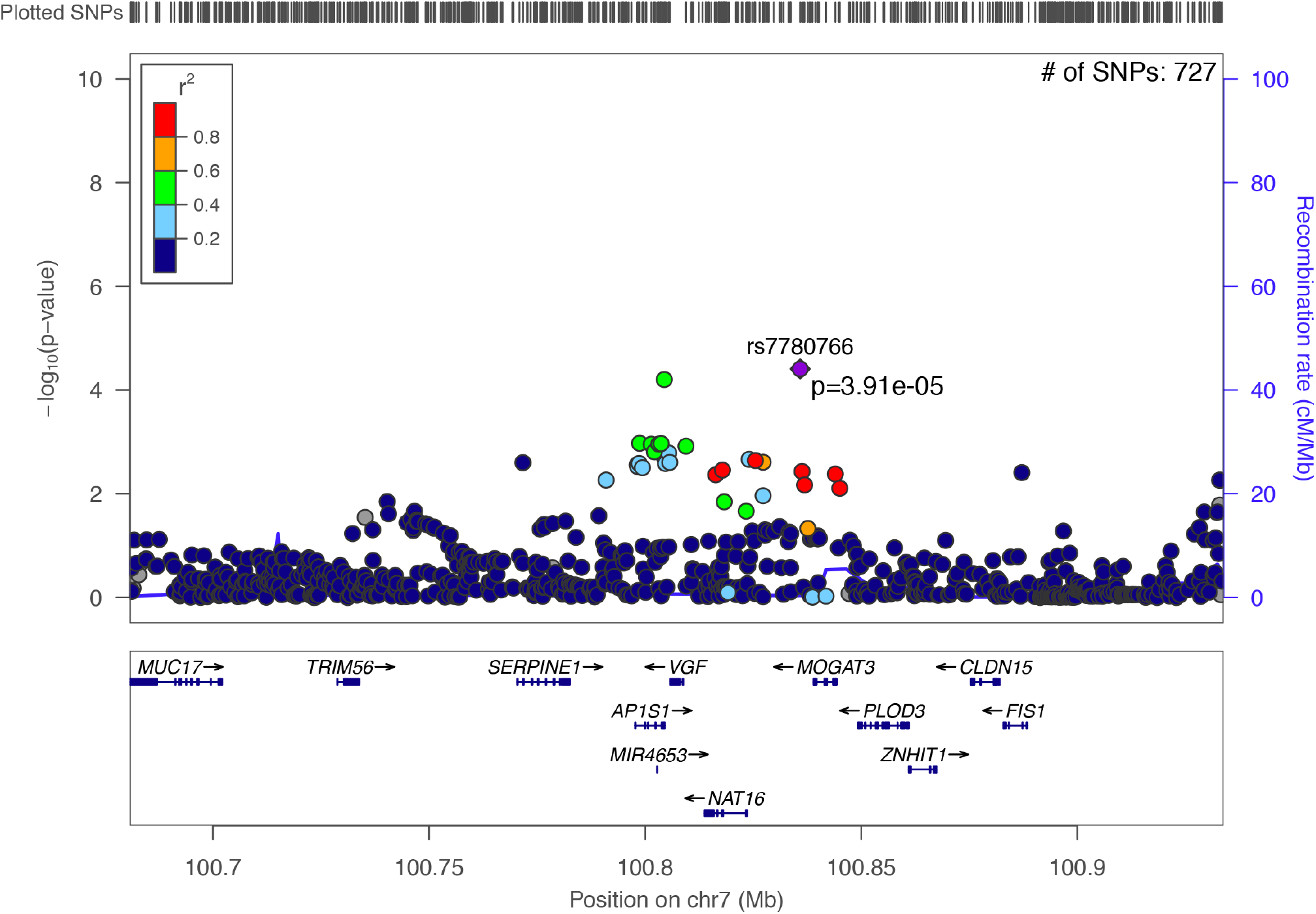
Genetic association of VGF to AD. Manhattan plot of the VGF locus in the I-GAP AD GWAS. SNPs in the locus show significant association to AD after Bonferroni correction.

**Fig. S6.**
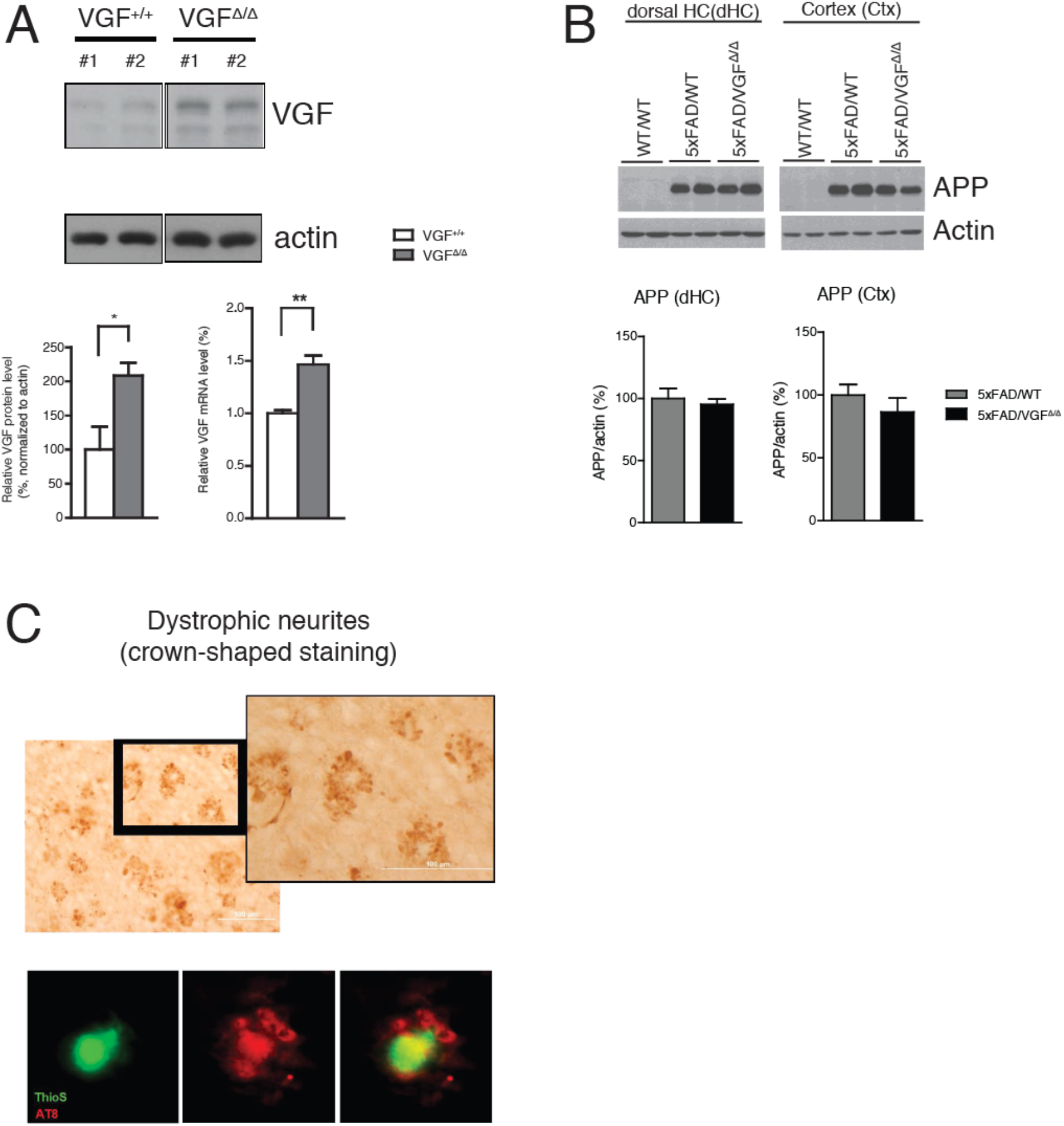
**A)** Increased VGF expression in the brains of VGF germline overexpression mouse line (VGF^Δ/Δ^). Western blot and quantitative PCR analysis showed both increased VGF protein and mRNA level in the dorsal hippocampus of VGF germline overexpression mice (hippocampus VGF protein: VGF^Δ/Δ^: 208.6±18.4%; WT: 100.0±33.7%; hippocampus Vgf mRNA: VGF^Δ/Δ^: 146.6±8.4%; WT: 100.1±3.0%, Student t-test, *, p<0.05; **, p<0.01. **B)** Similar levels of transgenic APP protein in both cortex and dorsal hippocampus of 5xFAD mouse brain with VGF germline overexpression. **C)** Amyloid plaques were surrounded by dystrophic neurite clusters. Upper panel: immunohistochemical staining of phospho-Tau (AT8) showed dystrophic neurite clusters. Lower panel: green, amyloid plaque (Thioflavin S staining); red, phosphor-Tau (AT8).

**Fig. S7.**
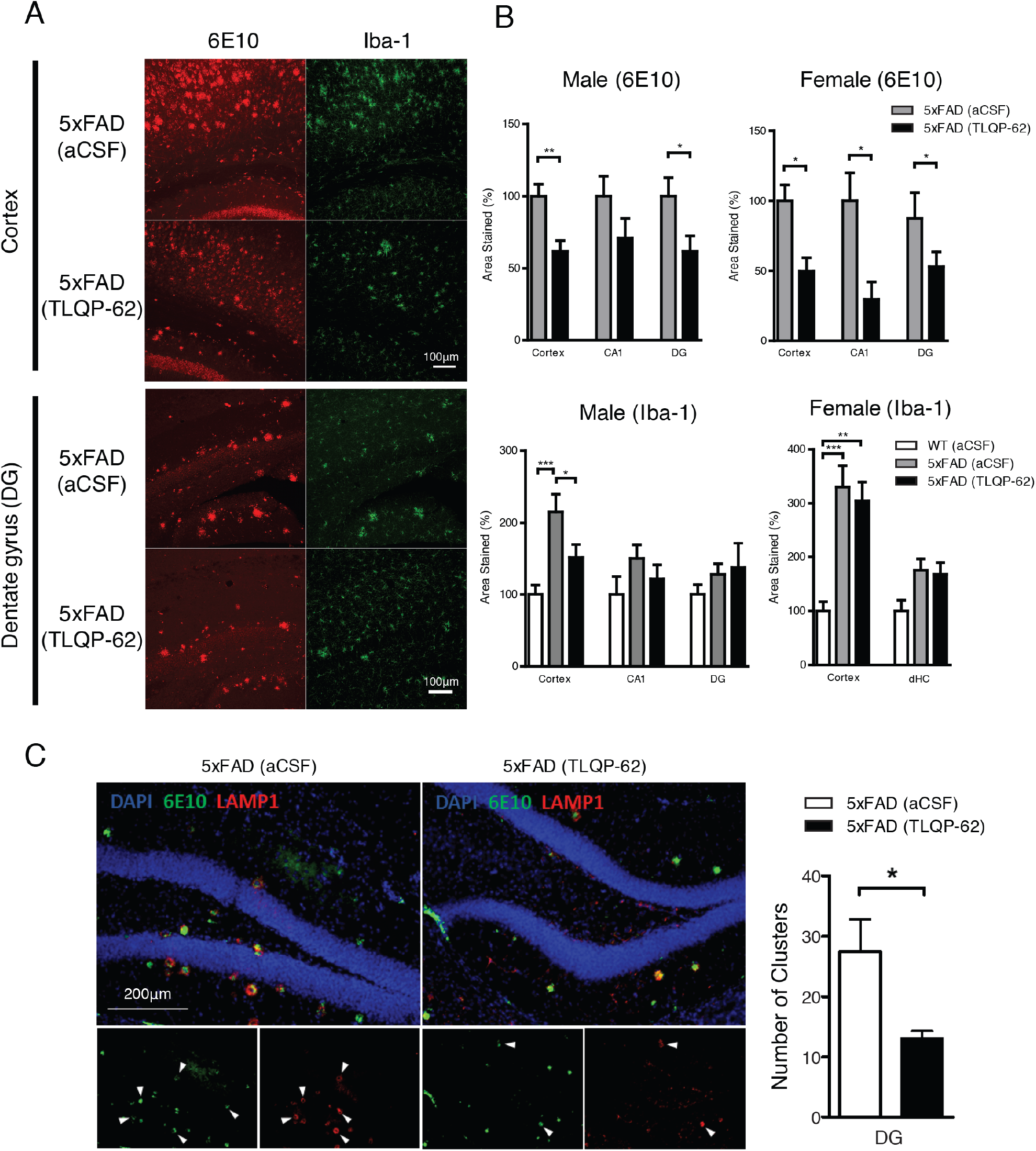
Chronic icv administration of TLQP-62 peptide ameliorated pathophysiological changes in the 5xFAD mouse brain. **A)** Immunohistochemical staining of Aβ amyloid plaques and microglial cells in the 5xFAD mouse cortex and dentate gyrus after 28-day icv administration of TLQP-62 peptide or vehicle control (aCSF). Red: Aβ (6E10), green: Iba-1. **B)** Quantification of percent area of Aβ and Iba-1 staining in both peptide-treated male and female 5xFAD mouse brains. N=4~5 mice/per group (male), 5~7 mice/per group (female). Data were analyzed by Student t-test (6E10 staining) or one-way ANOVA with Newman-Keuls post hoc analysis (Iba-1 staining). *: p<0.05, **: p<0.01, ***: p<0.001. **C)** Reduced staining of Lamp1-immunoreactive dystrophic neurite cluster number in 5xFAD brains after 28-day TLQP-62 icv infusion. Red: Lamp1, green: 6E10, blue: DAPI. N= 5~6 male mice/per group. Data were analyzed by Student t-test. *: p<0.05.

**Fig. S8.**
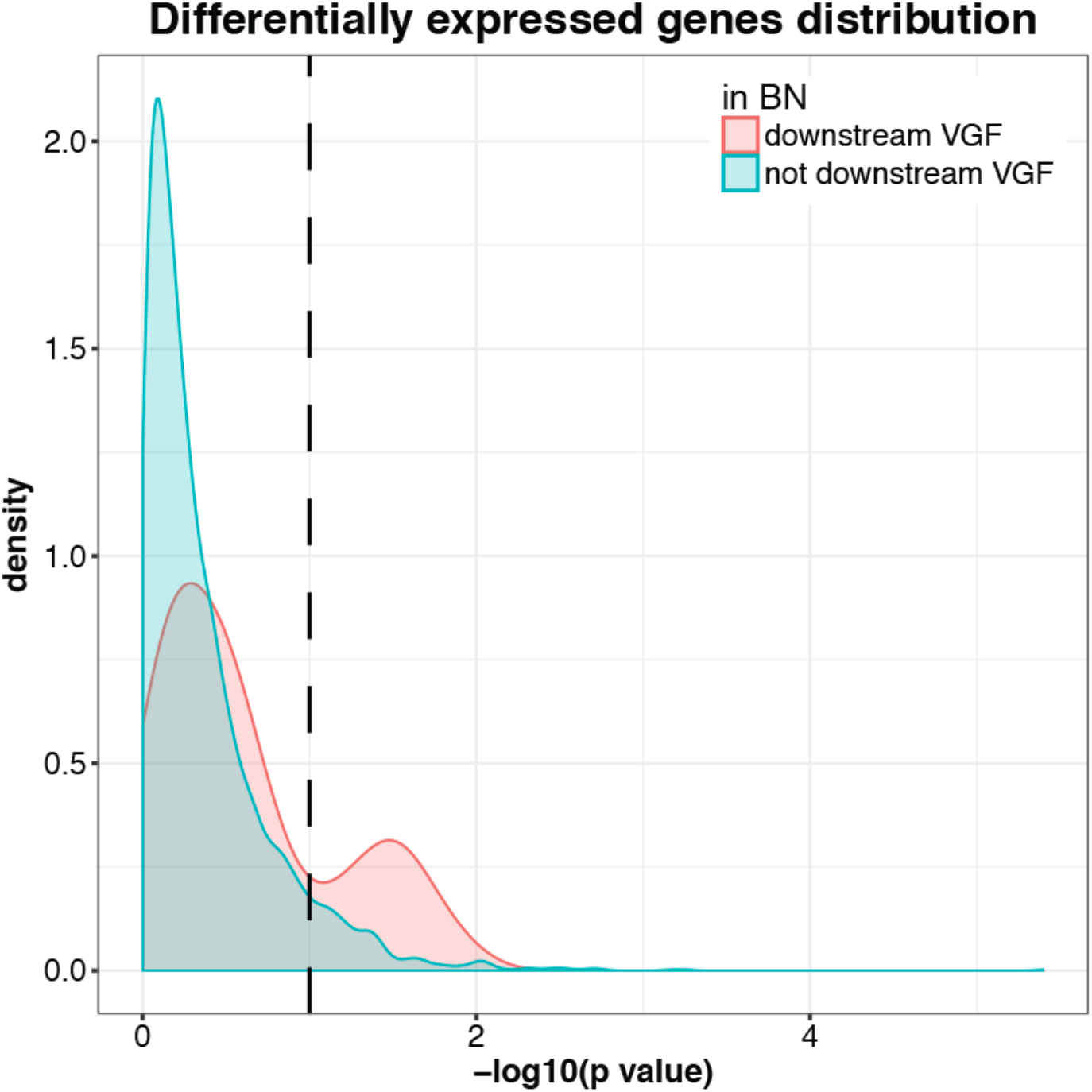
Density plot of the distribution of differential expression nominal p-values for genes downstream and not downstream (causally independent of the expression levels) of VGF in the gene-only network for mouse DE genes (5xFAD, WT versus 5xFAD, VGFΔ/Δ brains): The x axis is the – log10(p-value) and the y axis the densities. The red and blue curves are for genes downstream and not downstream of VGF in the network respectively.

**Table S1.** Classification of AD: This table defines the classification of samples in disease categories (152, 153) (for ROSMAP details, see https://www.synapse.org/#!Synapse:syn3191090).

| Dataset | classifier | controls | AD | definite controls | definite AD (dAD) |
| --- | --- | --- | --- | --- | --- |
| MSBB | PlaqueMean | continuous | continuous | < 6 | >= 12 |
| MSBB | CDR | < 1 | >= 1 | 0 | >= 1 |
| MSBB | CERJ | < 2 | >= 2 | 1 | 2 |
| MSBB | Path DX | controls | non-controls | controls | dAD |
| MSBB | bbscore | < 3 | >= 3 | < 3 | >= 3 |
| MSBB | NP-1 | < 2 | >= 2 | 1 | 2 |
| ROSMAP | Braaksc | < 3 | >= 3 | < 3 | >= 3 |
| ROSMAP | Ceradsc | >= 4 | < 4 | 4 | 1 |
| ROSMAP | Cogdx | < 4 | >= 4 | 1 | [4, 5] |

